# Clues to reaction specificity in PLP-dependent fold type I aminotransferases of monosaccharide biosynthesis

**DOI:** 10.1101/2021.09.04.459008

**Authors:** Jaya Srivastava, Petety V. Balaji

## Abstract

Novel functions can emerge in an enzyme family while conserving catalytic mechanism, motif or fold. PLP-dependent enzymes have evolved into seven fold-types and catalyse diverse reactions using the same mechanism for the formation of external aldimine. Nucleotide sugar aminotransferases (which will be henceforth referred to as aminotransferases) belong to fold type I and mediate the biosynthesis of several monosaccharides. They use diverse substrates but are highly selective to the C3 or C4 carbon to which amine group is transferred. Profile HMMs were able to identify aminotransferases but could not capture reaction specificity. A search for discriminating features led to the discovery of sequence motifs that are located near the pyranose binding site suggesting their role in imparting reaction specificity. Using a position weight matrix for this motif, we were able to assign reaction specificity to a large number of aminotransferases. Inferences from this analysis set way for future experiments that can shed light on mechanisms of functional diversification in nucleotide sugar aminotransferases of fold type I.

## 1. INTRODUCTION

Monosaccharides are the building blocks of carbohydrates and constitute the ‘third alphabet of life’. This alphabet has many more building blocks than even the protein alphabet^1^ and this is a consequence of glycodiversification^2,3^. Structural heterogeneity of monosaccharides is a functional requirement, especially among prokaryotes as they adapt to varying environmental conditions by evolving novel monosaccharides and glycans^2^. Glycan building blocks differ from each other primarily in the configuration and functional group substituents of carbon atoms^4,5^. These variations are brought about by only a handful of enzyme families as revealed by an analysis of the biosynthesis pathways of 55 monosaccharides^6^. These enzyme families are involved in other biological processes as well. For instance, a sugar 3-dehydrogenase that catalyses the biosynthesis of UDP-Man2NAc3NAcA^7^ belongs to the GFO/IDH/MocA family and this family contains enzymes of glycolysis and galactose metabolism also^8^. Clearly, organisms recruit existing enzymes, tweak substrate specificity and reaction mechanism through sequence and structural modifications and thereby generate novel enzymes.

The conserved core of an enzyme family can be defined by (i) a few residues, such as cofactor binding lysine in PLP dependent enzymes^9^, (ii) a motif, as that in Radical SAM superfamily^10^, or 3) a conserved fold, as that in Short chain dehydrogenase Reductases (SDRs)^11^ (Table 1). PLP-dependent enzymes use the same mechanism for the formation of external aldimine even though they catalyze different types of reactions^12^. They have been grouped into seven fold types based on a comparison of their 3D structures^13^ (Figure 1) thereby reflecting catalytic strategies that have independently evolved among PLP dependent enzymes^12^. Nucleotide sugar aminotransferases and dehydratases belong to fold type I^14^. This group also contains enzymes that catalyze several other types of reactions^15^ (Figure 1).

**Table 1:**
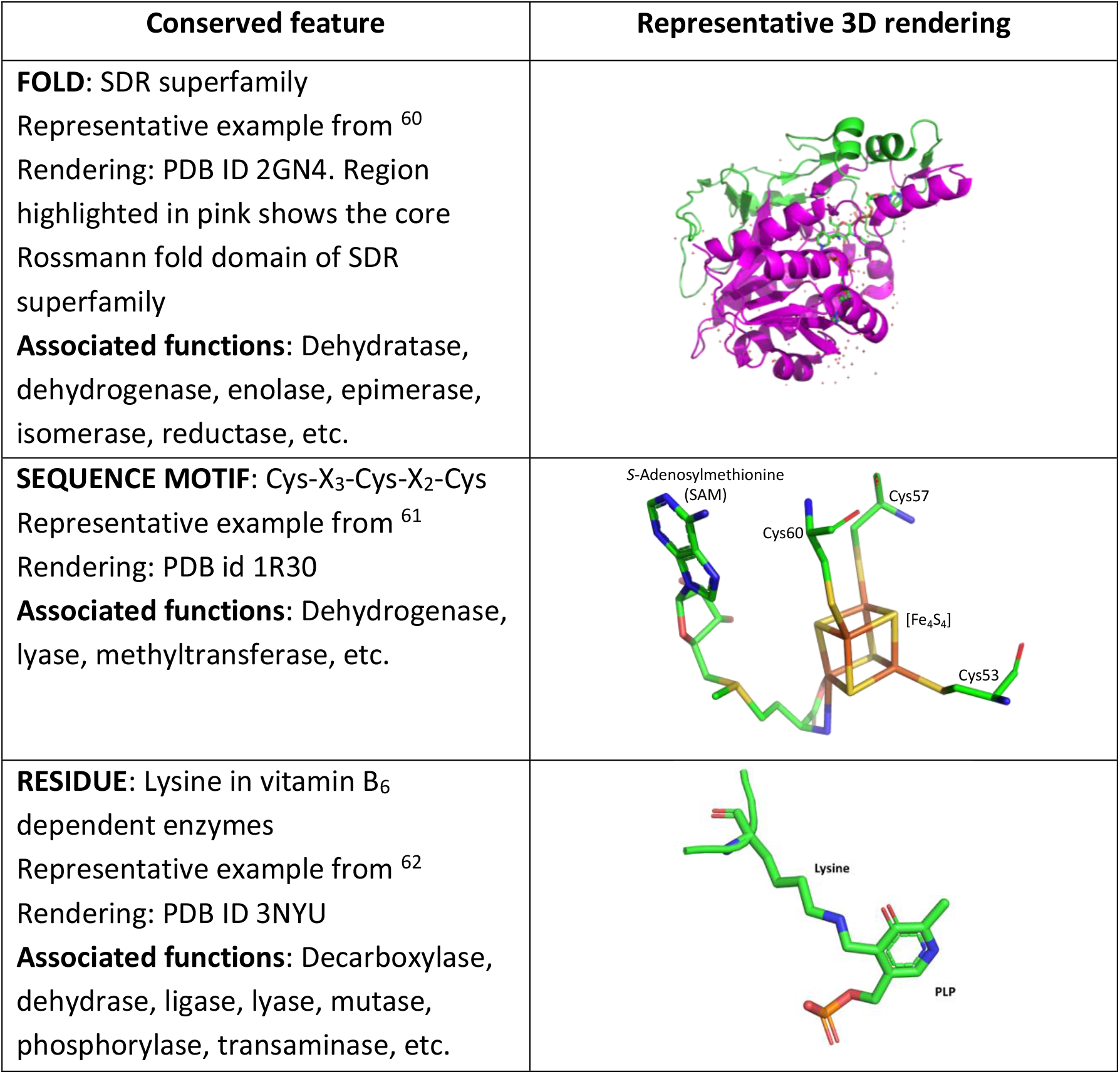
Features conserved at various levels among functionally diverse enzymes and illustrative examples

| Conserved feature | Representative 3D rendering |
| --- | --- |
| **FOLD:** SDR superfamily<br>Representative example from <sup>60</sup><br>Rendering: PDB ID 2GN4. Region highlighted in pink shows the core Rossmann fold domain of SDR superfamily<br>**Associated functions:** Dehydratase, dehydrogenase, enolase, epimerase, isomerase, reductase, etc. |  |
| **SEQUENCE MOTIF:** Cys-X<sub>3</sub>-Cys-X<sub>2</sub>-Cys<br>Representative example from <sup>61</sup><br>Rendering: PDB id 1R30<br>**Associated functions:** Dehydrogenase, lyase, methyltransferase, etc. |  |
| **RESIDUE:** Lysine in vitamin B<sub>6</sub> dependent enzymes<br>Representative example from <sup>62</sup><br>Rendering: PDB ID 3NYU<br>**Associated functions:** Decarboxylase, dehydrase, ligase, lyase, mutase, phosphorylase, transaminase, etc. |  |

**Figure 1:**
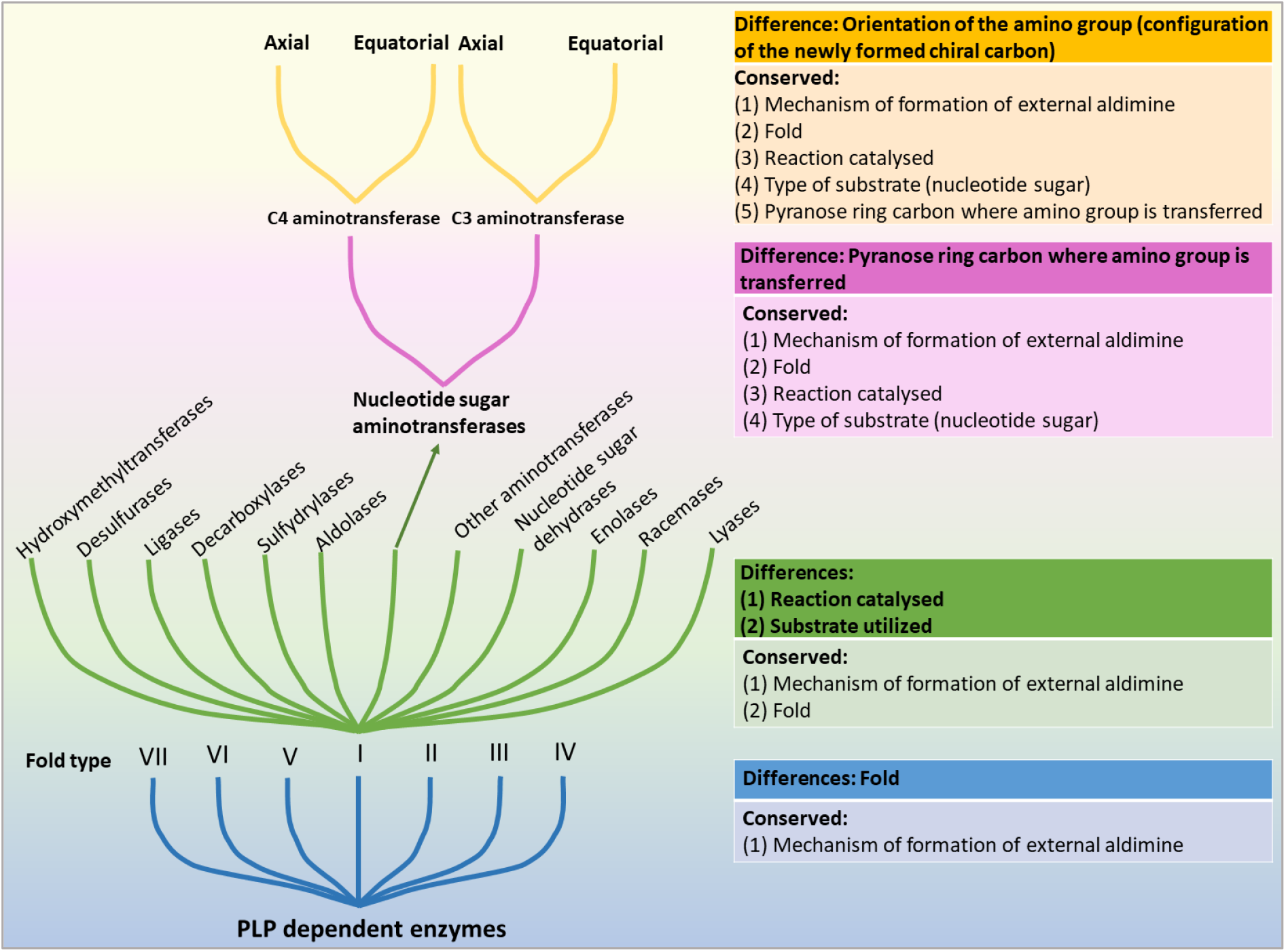
Neofunctionalization in PLP dependent enzymes leading to the evolution of specialized functions. Nucleotide sugar aminotransferases and dehydratases adopt fold type I. ‘Other aminotransferases’ include non-nucleotide linked aminotransferases as well as those utilizing distinct substrates such as amino-acids. Based on the pyranose ring carbon which is aminated, aminotransferases are further classified as C3 aminotransferases and C4 aminotransferases. These, in turn, can install amine either axially or equatorially (assuming ^4^*C*_1_ conformation) [Table S1, S2].

Aminotransferases and dehydratases take part in the biosynthesis of at least 19 monosaccharides (Figure S1) and secondary metabolites^16,17^. Aminotransferases convert keto-sugars to deoxy amino sugars. Dehydratases utilize similar substrates as aminotransferases but catalyse dehydration instead of transamination. For example, ColD (a dehydratase), utilizes the same substrate as Per (a C4 aminotransferase) but catalyzes dehydration as opposed to transamination^18^. These catalytic differences among aminotransferases and dehydratases have emerged by keeping the overall structure intact (Figure 2). Even the extent of sequence similarity among functional variants of aminotransferases and dehydratases is so comparable (Figure S2) that it is not possible to distinguish between these two families solely based on pairwise sequence similarity. Residues which upon mutation could convert dehydratase to aminotransferase have been reported in literature^19,20^. However, features governing reaction specificity among aminotransferases remain unexplored.

**Figure 2.**
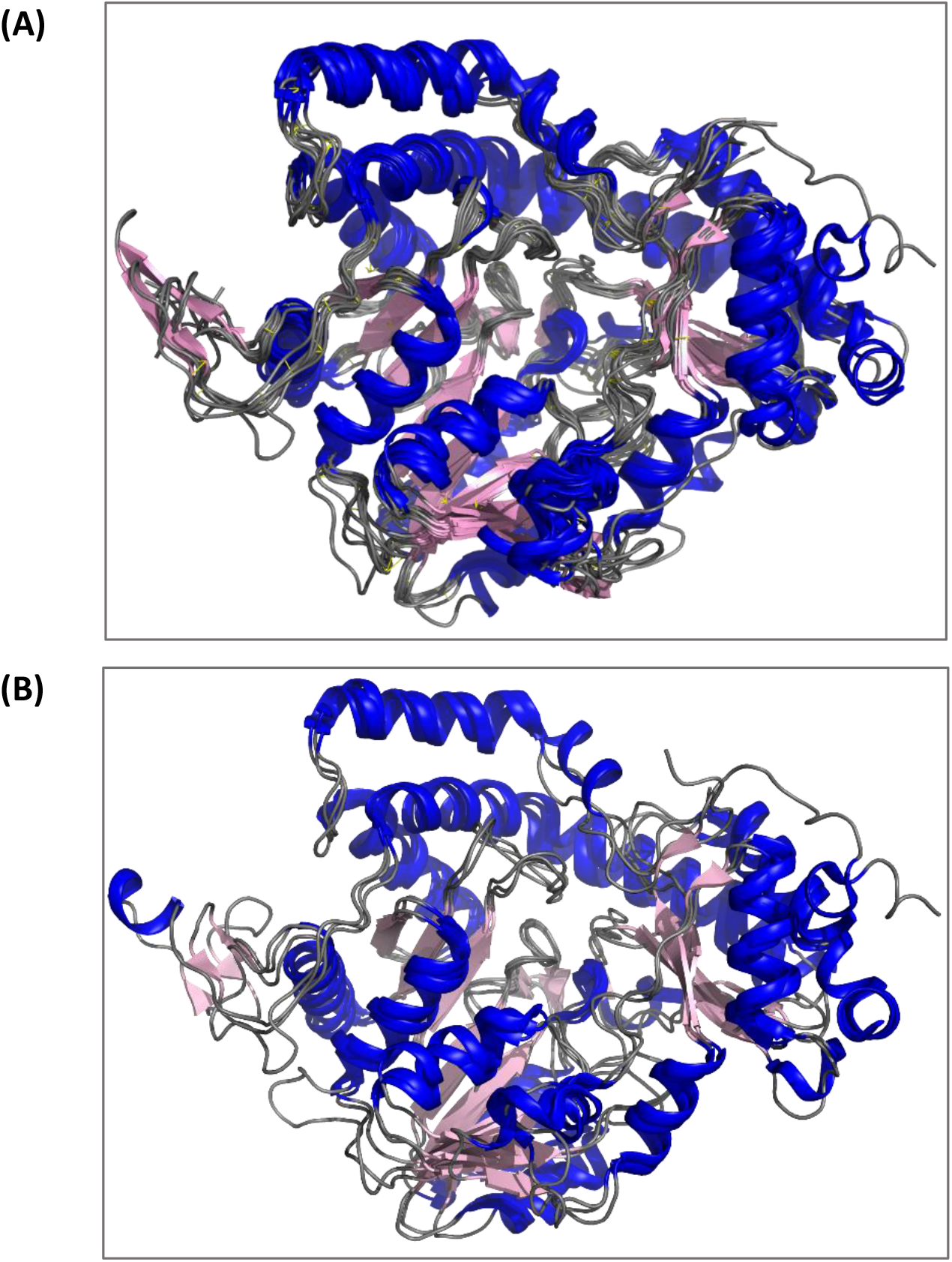
**A:** Structural superimposition of four C3 aminotransferases and six C4 aminotransferases. PDB ids, substrates and orientation of the amino group in the product are given below. **C3 aminotransferases** 3NYU^62^ UDP-2-acetamido-2-deoxy-3-oxo-glucuronate Equatorial (^4^*C*_1_) 5U21^62^ TDP-3-keto-6-deoxy-galactose/glucose Equatorial (^4^*C*_1_) 3FRK^24^ TDP-3-keto-6-deoxy-glucose Equatorial (^4^*C*_1_) 2OGA^63^ TDP-3-keto-4,6-dideoxy-D-*erythro*-hexopyranose Equatorial (^4^*C*_1_) **C4 aminotransferases** 4ZAH^64^ TDP-4-keto-6-deoxy-glucose Axial (^4^*C*_1_) 2PO3^42^ TDP-4-keto-6-deoxy-glucose Equatorial (^4^*C*_1_) 3DR4^65^ GDP-4-keto-6-deoxy-mannose Equatorial (^4^*C*_1_) 2FNU^66^ UDP-2-acetamido-2,6-dideoxy-beta-L-*arabino*-4-hexulose Equatorial (^4^*C*_1_) 1MDO^67^ UDP-beta-L-*threo*-pentapyranos-4-ulose Equatorial (^4^*C*_1_) 4ZTC^37^ UDP-4-keto-6-deoxy-GlcNAc Equatorial (^4^*C*_1_) **B:** Structural superimposition of a C3 aminotransferase (3NYT), a C4 aminotransferase (2PO3) and a dehydratase (2GMS). Other details (UniProt accession number, length, organism, gene / protein name and PubMed id) are given in Supplementary data.xslx, worksheet: Exp and reviewed sequences.

Aminotransferases may differ from each other in three ways: (i) substrate specificity, (ii) whether amino group is transferred to the C3 or C4 carbon of the pyranose ring, and (iii) the configuration of the chiral center formed as a consequence of amino group transfer (Figure 3). Some of the experimentally characterized aminotransferases (Supplementary data.xslx, worksheet: Exp and reviewed sequences) are specific to the nature and orientation of substituents on pyranose ring^21^ whereas others exhibit broad substrate specificity^22,23^. However, they are specific towards the position and orientation of amine group installation^21,23,24^. Clearly, reaction specificity is determined by as yet unknown “minor” differences in primary structures. In this study, we have identified features that discriminate C3 aminotransferases from C4 aminotransferases using profile Hidden Markov Models (HMMs) and position weight matrices (PWMs).

**Figure 3:**
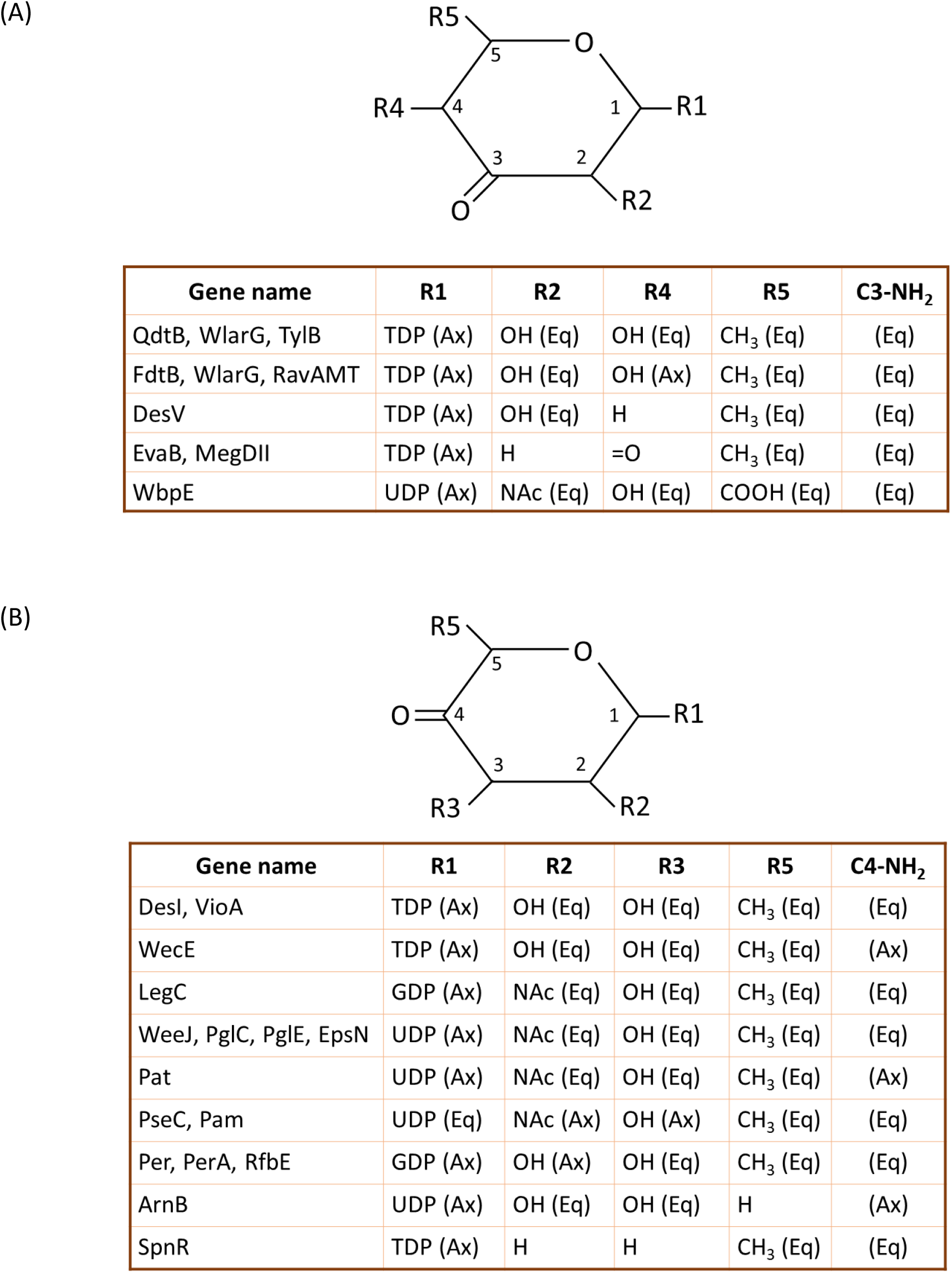
Composite figures showing the reactants and products of C3 aminotransferases (A) and C4 aminotransferases (B). Eq and Ax denote equatorial and axial, respectively. The orientation of the -NH_2_ group in the product is shown in the right-most column. Ax (axial) and Eq (equatorial) are included in parenthesis merely to bring out the similarities in the orientations of the substituents on the pyranose rings in the substrates of various enzymes; here, the ^4^*C*_1_ conformation is assumed for the pyranose ring. Schematic structures of reactants and products are given in Tables S1, S2.

## 2. METHODS

### Databases, software and online servers

Protein sequences were obtained from UniProtKB (Release 2021_1)^25^ and 3D structures from the Protein Data Bank^26^. Sequences which had one or more of the characters B, O, J, U, X and Z were ignored. The CATH database (v4.2) was used for identifying proteins which share the same fold^27^. Pairwise sequence comparisons were performed using BLASTp online server^28^. Pairwise identity matrix was obtained using MUSCLE webserver^29^. Multiple sequence alignments (MSAs) were done using the MAFFT command line application^30^. 3D structures were visualized, analyzed and compared using PyMol^31^. HMMER3^32^ was used to build profile HMMs. Electron density maps of ligand structures were visualized using WinCOOT^33^.

### Datasets and profile HMMs

Experimentally characterized aminotransferases and dehydratases were obtained from literature; reviewed homologs were taken from SwissProt (Supplementary data.xslx, worksheet: Exp and reviewed sequences). Bit score thresholds for NSAT_HMM_ and NSD_HMM_, profile HMMs of aminotransferases and dehydratases, respectively, were set based on Receiver Operator Characteristic (ROC) curves generated using hits from TrEMBL under the assumption that annotations provided in TrEMBL are correct. The procedure used to generate ROC curves is described in detail elsewhere^6,34^. Briefly, ROC curves were generated by calculating the number of true positives, false positives and false negatives for various values of bit score threshold (Figure S3). TrEMBL entries which did not have molecular function annotation were ignored. Possible thresholds chosen from the inflection point of ROC curve were validated by scoring the experimentally characterized sequences of CATH superfamily 3.40.640.10 (fold type I PLP-dependent enzymes) [Supplementary data.xslx, worksheet: CATH-3.40.640.10]. Based on this procedure, thresholds for NSAT_HMM_ and NSD_HMM_ were set to be 310 and 300 bits, respectively.

ROC curves could not be used to set thresholds for C3_NSAT_HMM_ and C4_NSAT_HMM_ (profile HMMs for C3 and C4 aminotransferases, respectively) since a majority of annotations in TrEMBL just mention aminotransferase without specifying if the protein is a C3 aminotransferase or C4 aminotransferase. To overcome this hurdle, an alternative approach was used whereby the TrEMBL database was searched with these two profiles separately by using the default value suggested by the HMMER software as threshold (viz., E-value = 10). A bit score scatter plot of hits common to both profiles was obtained and on the basis of this plot, thresholds for both C3_NSAT_HMM_ and C4_NSAT_HMM_ were set to 400 bits (Figure 4A).

**Figure 4:**
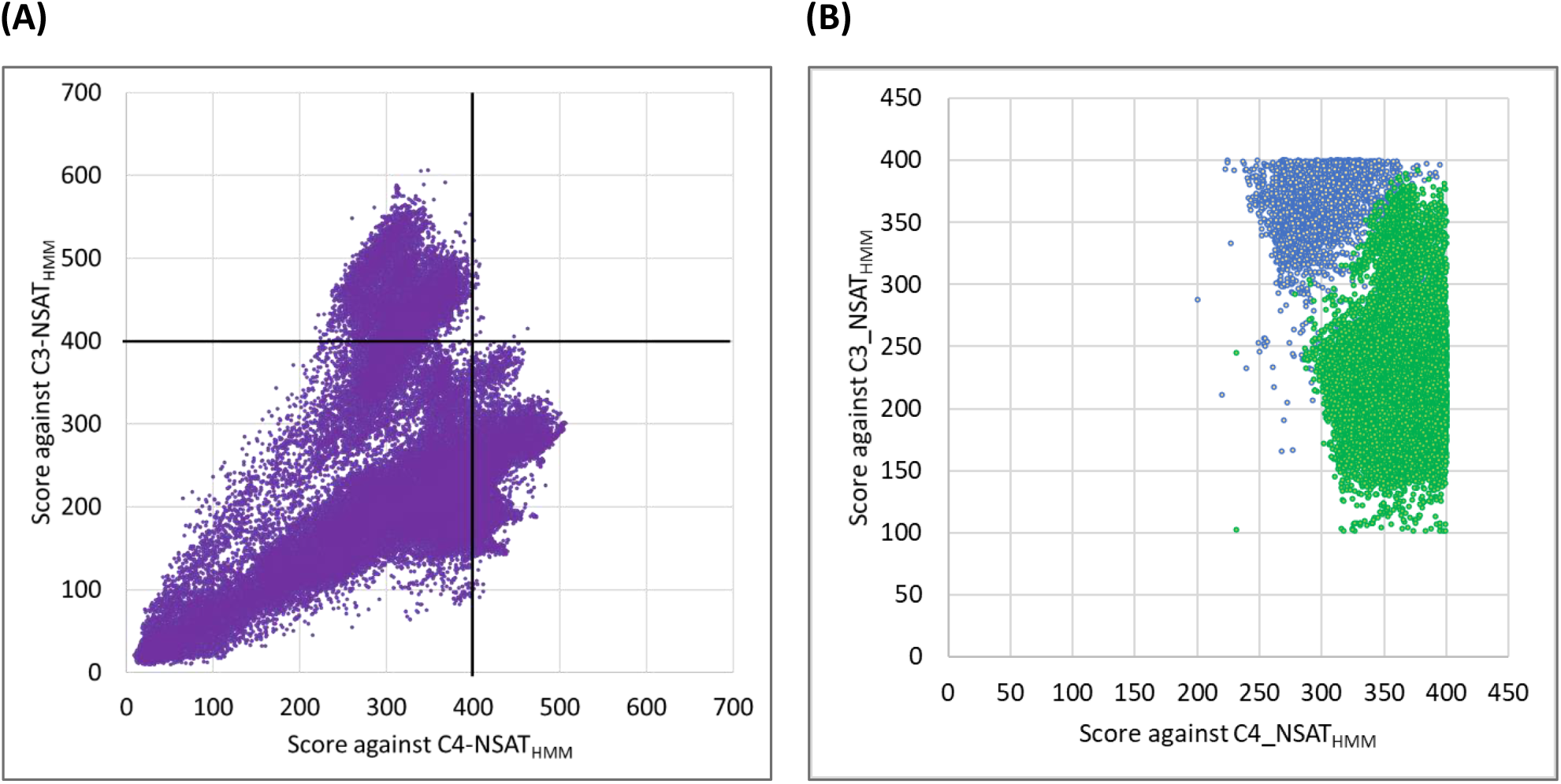
(A) Distribution of bit scores of hits common to both C3_NSAT_HMM_ and C4_NSAT_HMM_ from TrEMBL using default threshold suggested by the HMMER software. As can be seen, some proteins are “high-scoring” for one profile while being “low-scoring” for the other profile. This formed the basis on which the threshold was set 400 bits for both the profiles (B) Comparison of bit score and PWM log-odds score prediction. Green dots represent sequences that are predicted as C4 aminotransferases by motif search, while blue dots are predicted as C3. Note: this plot does not include 1,322 sequences for which PWM-based prediction was inconclusive.

### Identification of sequence motifs

The command line application of STREME^35^ was used to identify sequence motifs enriched in a sequence family with reference to a set of “control” sequences. By subjecting TrEMBL hits of C3_NSAT_HMM_ and C4_NSAT_HMM_ to a redundancy criterion of 100% identity, a total of 12,811 and 12,101 sequences were obtained and designated as C3_hits and C4_hits, respectively. The dataset C4_hits was used as control sequence dataset to discover motifs in C3_hits and vice versa. Position-specific amino acid frequency matrices were obtained from STREME and converted into a PWM of log odds score viz., 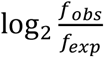 where *f_obs_* and *f_exp_* are observed and expected amino acid frequencies in a given position of the MSA. Expected frequencies were fetched from TrEMBL (Release 2021_1). For a given sequence, the region that has the highest log odds score was assumed to be the best possible match for the motif; in fact, the highest scoring motifs were verified to be the motifs of interest using in-house python scripts. Steps leading to motif discovery using profile HMMs are schematically represented in Figure 5A.

**Figure 5:**
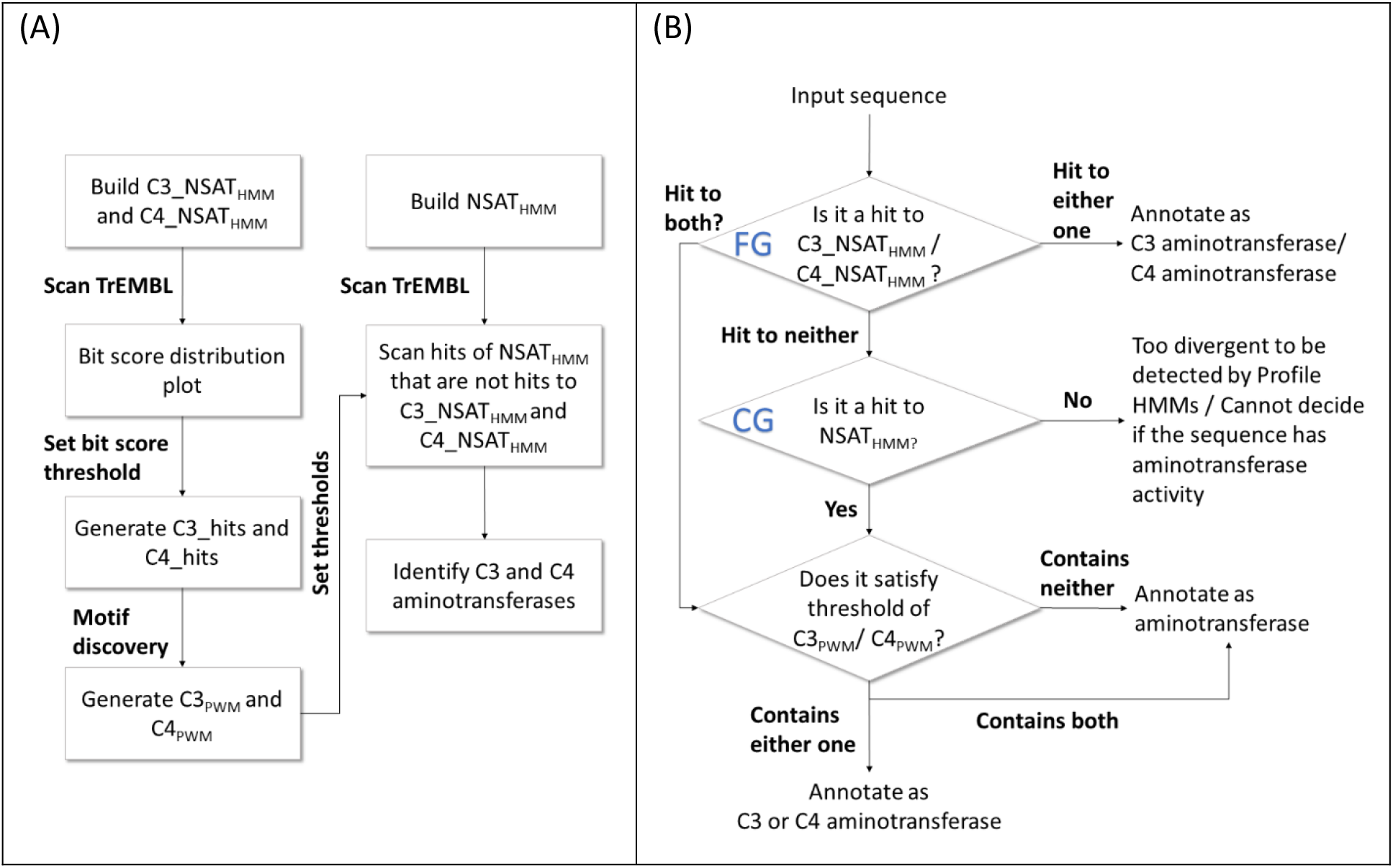
(A) Steps involved in identification of C3 and C4 aminotransferases from TrEMBL using profiles HMMs and PWMs. (B) Flowchart showing the steps involved in annotating an input sequence based on profile HMMs and PWMs. In steps marked FG and CG, annotations are Fine Grained, i.e., C3 aminotransferases an C4 aminotransferases, and Coarse Grained, i.e., aminotransferases (relative to each other; also see Figure 1).

### Phylogenetic analysis

Phylogenetic analysis was performed using MEGA7; sequences were first aligned by MUSCLE using default values for all the parameters and then creating a phylogenetic tree by Maximum Likelihood method^36^. Experimentally characterized sequences from CATH superfamily 3.40.640.10 (Supplementary data.xslx, worksheet: CATH-3.40.640.10) with >=80% identity were removed, and the resulting 208 enzymes, including PglE from *Camplyobacter jejuni* (PDB ID: 4ZTC)^37^, a C4 aminotransferase that is structurally identical to other C4 aminotransferases that is yet to be updated in CATH, were used for phylogenetic analysis. The tree was visualized using iTOL^38^.

## 3. RESULTS

### Identifying homologs of aminotransferases using profile HMMs

Scoring TrEMBL using C3_NSAT_HMM_ and C4_NSAT_HMM_ resulted in ~14000 hits for each profile. Twenty-one hits satisfied the threshold of both the profiles (viz., 400 bits) (Supplementary data.xslx, worksheet: common_hits). Among sequences scoring below 400 against both profiles, 32,344 were hits to NSAT_HMM_. These sequences are predicted to be aminotransferases (because they are hits of NSAT_HMM_) but we could not resolve if these are C3 or C4 aminotransferases, or neither of these two. Similarly, we could not resolve if 21 common hits are C3 or C4 aminotransferases or have both activities. Thus, our inability to set a bit score threshold that is sensitive enough to identify C3 and C4 aminotransferases without compromising specificity meant that an alternative approach is required to discriminate C3 and C4 aminotransferases from each other.

### Identifying motifs that discriminate C3 aminotransferases and C4 aminotransferases from each other

C3_hits and C4_hits were used to find motifs enriched in C3 aminotransferases in comparison to C4 aminotransferases and vice versa using STREME^a^. The most conserved motif among C3_hits is a 15-mer found in 99.9% of input sequences (Figure 6A,6B). The most conserved motif among C4 aminotransferases is a 12-mer located at the same site as the conserved motif in C3_hits (Figure 6A,6C). This motif is found in all but one C4_exp sequences. Visual inspection of the sequence logos confirmed the distinctiveness of these motifs (Figure 6A). Specifically, the conserved 15-mer of C3_hits contain a glycine residue at position 9 (Figure 6B), whose role has not been investigated so far. Both motifs map to the active site of aminotransferases (Figure 6D). We postulate that these motifs influence the positioning of the substrate in the binding site for C3 or C4 amine installation. These motifs are henceforth referred to as C3_motif and C4_motif, respectively.

**Figure 6:**
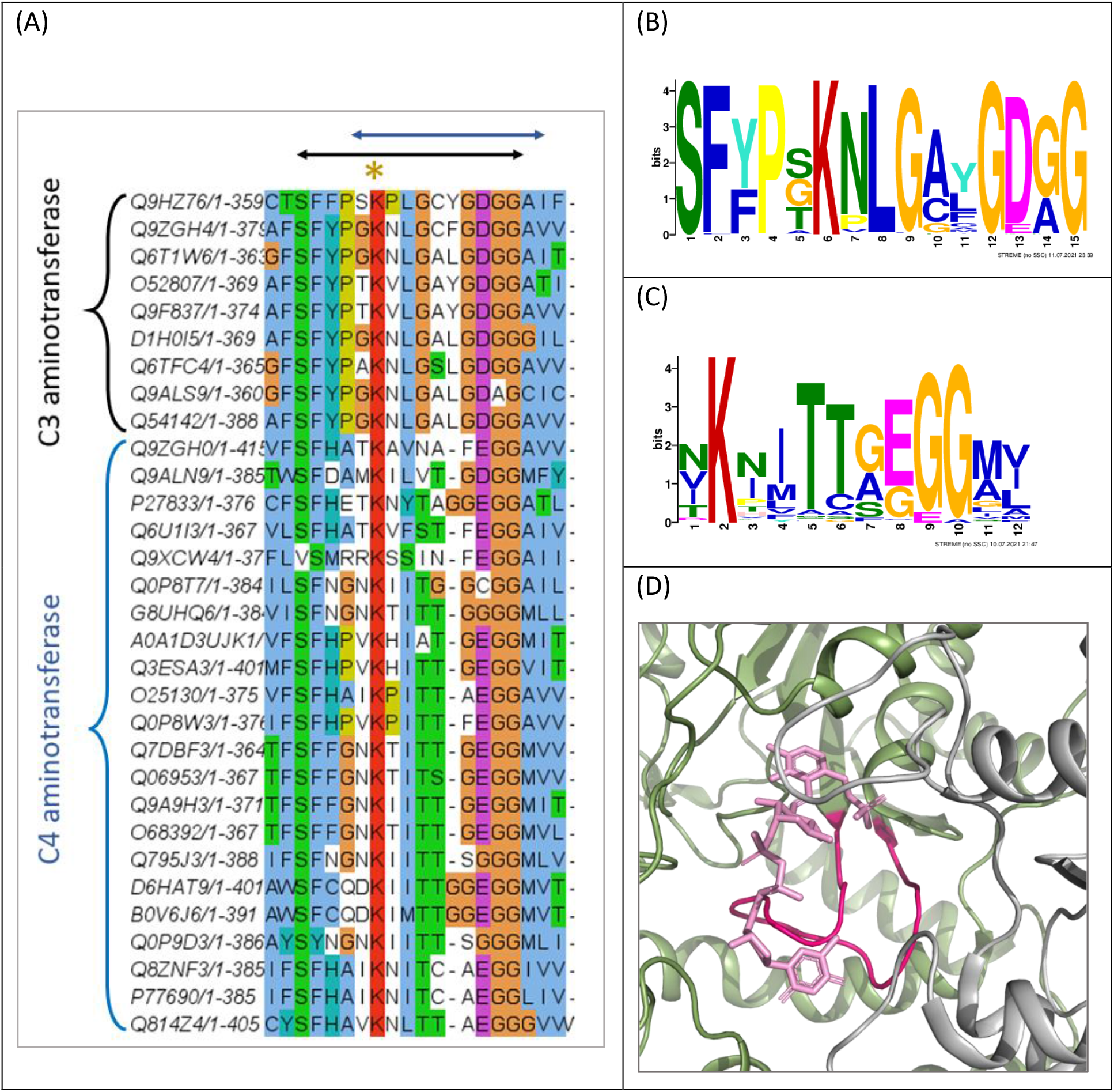
**(A)** A part of the multiple sequence alignment of C3 and C4 aminotransferases. The conserved lysine that serves the role of Schiff’s base is marked with a brown asterisk. The 15-residue (in C3 aminotransferases) and 12-residue (in C4 aminotransferases) motifs identified by the STREME server are marked by black and blue double-headed arrows, respectively. Whether a 12-mer corresponding to the 12-mer of C4 aminotransferases can be a potential C3_motif was also probed. It was found that the highest scoring 12-mers in six sequences from C3_hits were not from the expected position, but adjacent to it. Hence, this 12-mer motif was not considered for any further analysis. Sequence logos of motifs obtained from STREME for C3_hits **(B)** and C4_hits **(C)**. **(D)** The location of the motif in the 3D structure of a C3 aminotransferase viz., QdtB from *Thermoanaerobacterium thermosaccharolyticum* (PDB ID 3FRK)^24^. The motif is in dark pink and the bound nucleotide sugar product is in light pink.

### Setting bit score threshold for PWMs

Two PWMs viz., C3_PWM_ and C4_PWM_, were generated using the MSAs of C3_motif and C4_motif, respectively. The entire length of each and every sequence in C3_hits and C4_hits was scanned by both these PWMs. The highest scoring 15-and 12-mer in each sequence was deemed to be the region that best matches the motif. The score of the highest scoring 15-/12-mer of a sequence was assigned as the score for that sequence. The distribution of scores showed the near-absence of 15-/12-mers that have positive scores when C3_hits are scanned using C4_PWM_ and when C4_hits are scanned using C3_PWM_ (Figure 7).

**Figure 7:**
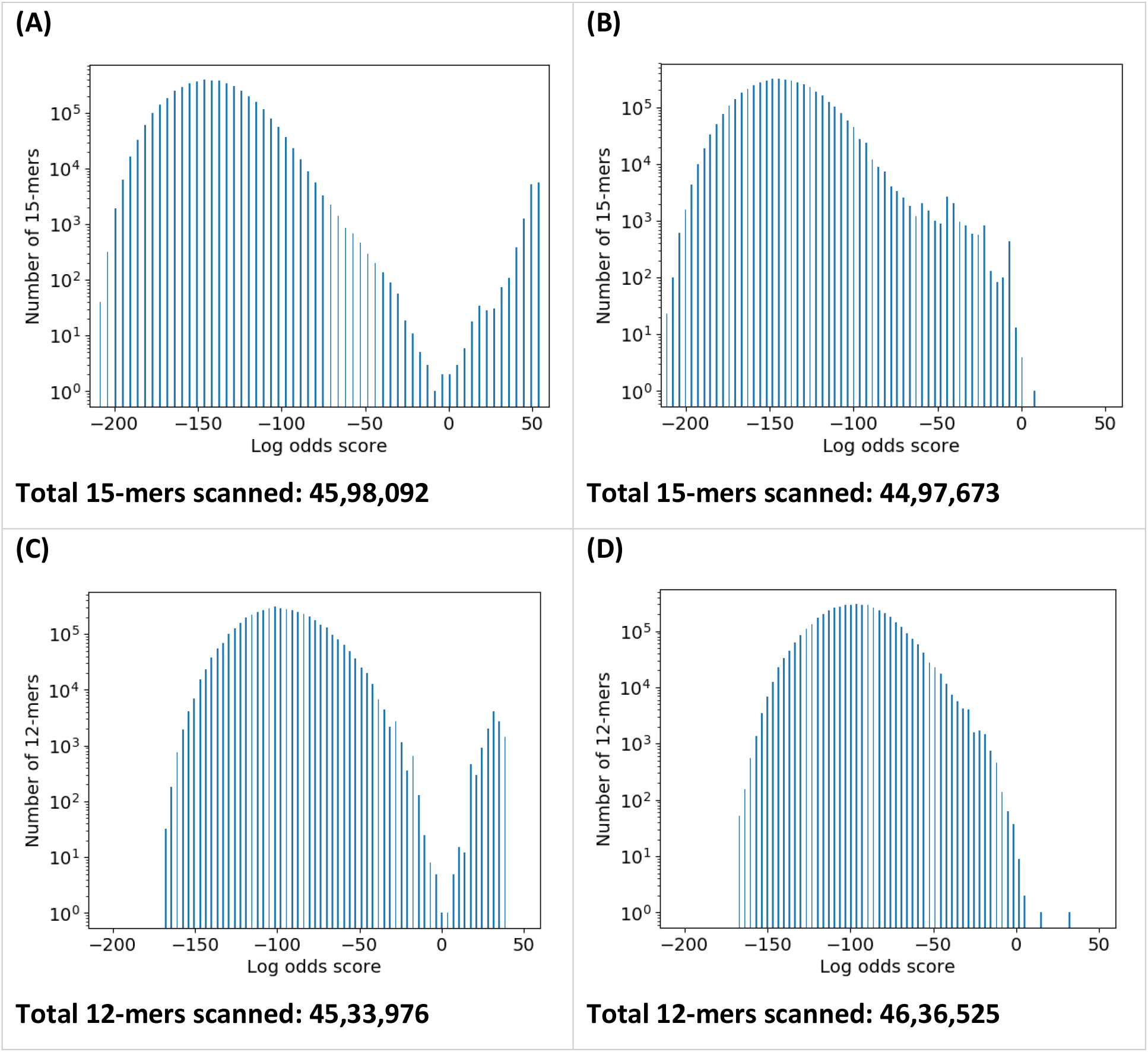
Distribution of log odds score of all 15-mers of C3_hits (A) and C4_hits (B) for the C3_PWM_ and all 12-mers of C4_hits (C) and C3_hits (D) for the C4_PWM_. Log-odds score is plotted in the X-axis and the number of 15-/12-mers (in log scale) are plotted in the Y-axis.

When C3_hits were scanned using C3_PWM_, four sequences scored <0 bits; three of these contain deletions in the C3_motif, including deletion of active site lysine; the fourth appears to be a C4 aminotransferase based on the high score (35 bits) when scanned against C4_motif. When C4_hits are scanned using C3_PWM_, five sequences score >0 bits; four among these score >30 bits when scanned against C4_PWM_ and the identity of the fifth sequence could not be ascertained based on C3_PWM_ or C4_PWM_.

When C4_hits are scanned using C4_PWM_, all the sequences score >6 bits except for one which scores 0.07 bits, because of a two-residue insertion. However, when C3_hits are scanned using C4_PWM_, 32 sequences score > 0 bits. One among these appears to be a C4 aminotransferase as it scores <0 bits against C3_PWM_. Three others score in the range 6-15 bits against C4_PWM_ but > 35 bits against C3_PWM_; their identities could not be resolved using motif search. The rest score in the range 0-3.5 bits against C4_PWM_ but > 30 bits against C3_PWM_. Based on these data, thresholds for C3_PWM_ and C4_PWM_ were set to 3 and 4 bits, respectively. Hence, with four exceptions, motif search captures differences among C3 aminotransferases and C4 aminotransferases that could not be identified using profile HMMs.

### Resolving common hits

As mentioned earlier, 21 sequences are hits to both C3_NSAT_HMM_ and C4_NSAT_HMM_ (Supplementary data.xslx, worksheet: common_hits). They were scanned using C3_PWM_ and C4_PWM_. 19 sequences are distinctly identified as C3 aminotransferase or C4 aminotransferase based on PWM scores. However, two proteins with accession numbers A0A2G9NIF3 and A0A1W9P6M8 satisfy the log-odds threshold of both motifs. Both profile HMM and PWM-based searches are not effective in resolving the specificities of these two proteins. Is it possible that these two enzymes have broad reaction specificity?

### Sorting out C3 aminotransferases and C4 aminotransferases among hits of NSAT_HMM_

Data from the scanning of C3_hits and C4_hits using the two PWMs show that C3_motif and C4_motif are unique to respective families of proteins. Hence, a two-step screening wherein the amino acid sequence of a protein is (i) first scored using C3_NSAT_HMM_ and C4_NSAT_HMM_, and (ii) then by C3_PWM_ and C4_PWM_ is proposed for the functional annotation of C3 aminotransferases and C4 aminotransferases (Figure 5B). The 32,344 sequences that score below the thresholds of C3_NSAT_HMM_ and C4_NSAT_HMM_ but are hits to NSAT_HMM_ were analyzed for the presence of C3_motif and C4_motif. 25,135 were predicted as C4 aminotransferases and 5,887 as C3 aminotransferases. It can be observed that C3 aminotransferases score less than C4 aminotransferases against C4_NSAT_HMM_; however, the converse is not true i.e., C4 aminotransferases do not necessarily score less than C3 aminotransferases against C3_NSAT_HMM_ (Figure 4B). 1073 and 249 sequences, respectively, score less than and higher than the thresholds of both PWMs. We are unable to identify their function based on the two-step process flow proposed herein. One possible reason is that the process is not sensitive enough. Another possibility is that some of these sequences have broad reaction specificity. Alternatively, these sequences may have lost aminotransferase activity and acquired novel function(s) due to sequence changes that are too subtle to be detected by HMMs and PWMs.

### Validating the prediction pipeline with putative aminotransferases of known biosynthesis pathways

We identified incompletely characterized sugar biosynthesis pathways from literature that involve putative aminotransferases to validate our two-step prediction pipeline. Seven such aminotransferases were identified (Supplementary data.xslx, worksheet: Test dataset). Four among these were identified as C3 aminotransferases using profile HMMs. The rest, OleNI, KijD7, and EryCIV, were identified as C4 aminotransferases using profile HMM and PWM-based search. OleNI was earlier proposed to act as a 3,4-dehydratase in desosamine biosynthesis^39^. However, the pathway for TDP-desosamine has been revised using homologous genes from *Streptomyces venezuelae*^40^ and *Saccharopolyspora erythraea*^41^ and supports our prediction of OleNI as a C4 aminotransferase.

### Additional factors governing product identity - clues from 3D structures

We superposed the 3D structures of C3 and C4 aminotransferases bound to substrates in the form of external aldimine (Table 2). The backbone conformation is highly similar in these six proteins (Figure 2A). In addition, the overall binding modes and conformations of ligands are also similar as can be seen from the overlap in the position of PLPs and of the nucleotides in the six complexes (Figure S4A). However, PLP forms aldimine with C3 keto or C4 keto group and this difference is accommodated by conformation change around the pyrophosphate group (Figure S4A). The conformation of ligands is identical in C3 aminotransferases (Figure S4B); in the three C4 aminotransferases, the pyranose ring either flips or shifts (Figure S4C, S4D). A tyrosine residue H-bonds with the N-acetyl / -OH group at C2 in all three crystal structures and this residue is conserved in C3_hits (Table 2). No such H-bond is observed in the three crystal structures of C4 aminotransferases, and this may be the reason for the observed shift/flip of the pyranose ring. Multiple sequence alignment of C4_hits shows that the homologous position is a tyrosine in ~40% of the sequences; in rest of the sequences, it is a tryptophan or phenylalanine. It has been proposed that ring flipping leads to variations in stereochemical outcome at C4^42^. Experimental verification is required to confirm the role of these surrounding residues as well as the discriminating sequence motifs that were used for PWM-based motif search, in determining the position of amine installation.

**Table 2:**
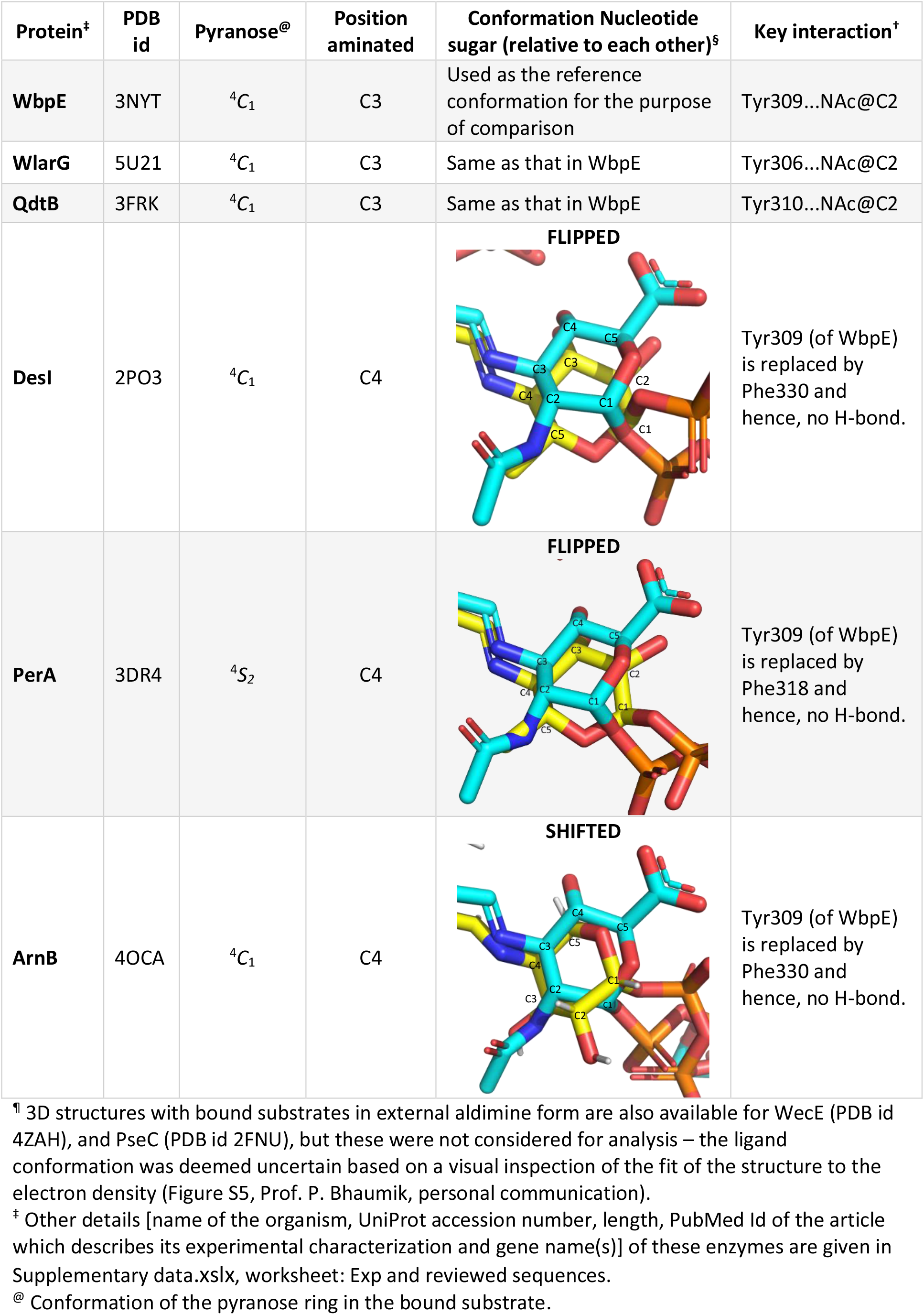
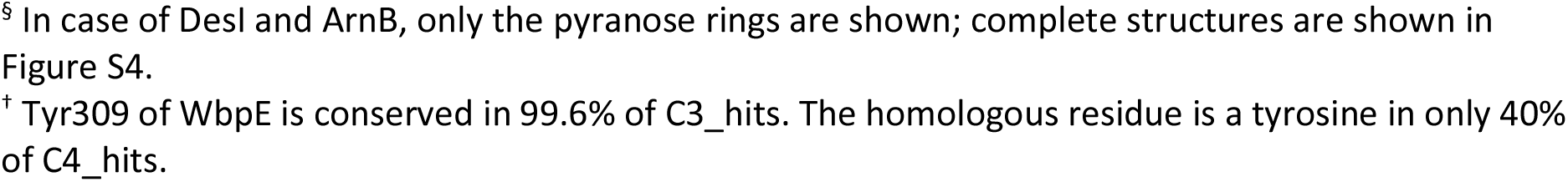
Aminotransferases with bound substrates in the form of external aldimine used for structural analysis^¶^

## 4. Discussion

Fold type I PLP-dependent enzymes constitute a family of enzymes wherein sequence and structural variations have led to functional diversification within a conserved fold (Figure 1). An earlier 3D structure-based phylogenetic study of fold type I PLP-dependent enzymes revealed a ‘sugar AT family’ consisting of aminotransferases, dehydratases, non-nucleotide sugar aminotransferases and AHBA synthase^43^. We obtained a similar sugar AT family of 22 enzymes from a sequence based phylogenetic analysis (Figure S6). Functions associated with other clusters also are largely conserved within a clade. The similarities in the 3D structure-based and sequence-based trees implies that sequence changes that lead to functional changes are reflected in 3D structures despite of fold conservation. What features govern reaction selectivity between C3 and C4 aminotransferases and between aminotransferases and dehydratases?

Aside from sequence and structural similarity among aminotransferases, conserved PLP and nucleotide group bound conformations suggest that differences are confined to regions that interact with pyranose ring (Figure S4). Using a combination of profile HMMs and motif based PWMs, we located a sequence motif in the pyranose binding site that differs among C3 and C4 aminotransferases (Figure 6). Structural comparisons led to the identification of a tyrosine residue that forms H-bond with pyranose ring and is conserved among C3 aminotransferases but not in C4 aminotransferases (Table 2). This suggests that absence of stabilizing interactions may lead to flipping or shifting of pyranose ring to orient C4 keto group for amination. H-bonding residues in that position are conserved among C3_hits (as tyrosine) but only in ~40% of the C4_hits. Thus, tyrosine can be one of the factors that determine the shift or flip observed in pyranose ring.

Other enzymes in the ‘sugar AT family’ contain dehydratases that utilize nucleotide linked sugars but catalyse dehydration instead. The overall reaction involves C3 deoxygenation, and is achieved by a single enzyme in the pathways for the biosynthesis of GDP-L-colitose and TDP-forosamine^18,44^ but by two enzymes in the pathway for the biosynthesis of CDP-linked 3,6-dideoxy sugars^19^. A hallmark of dehydratases is the replacement of active site lysine with histidine, though its replacement with lysine does not convert it to an aminotransferase, but merely stalls its native activity ^20^. Additional residues, i.e., Asp194, Tyr217 and Phe345 in *Yersinia pseudotuberculosis* E1 dehydrase and Ser187 in *Escherichia coli* ColD, have been shown to convert a dehydratase to an aminotransferase ^19,20^. However, among experimentally characterized dehydratases (Supplementary data.xslx, worksheet: Exp and reviewed sequences), Ser187 is not conserved, D194 is also found substituted with glutamate which is found among several C4 aminotransferases, and Y217 is conserved among C3 aminotransferases as well. We sought to analyse the extent of conservation in these sites, i.e., positions corresponding to Asp194 (Site 1), Tyr217 (Site 2) His220 (site 4), and Phe345 (Site 5) (*Yersinia pseudotuberculosis* E1 dehydratase numbering) and Ser187 (Site 3) (*Escherichia coli* ColD numbering) among NSD_hits (a dataset of 4018 TrEMBL entries that satisfy the bit score threshold of NSD_HMM_), C3_hits and C4_hits (Table S3). Observations from this analysis suggest that not all these sites are strictly conserved among dehydratases. ‘Sugar AT family’ also contains aminotransferases that utilize sugars not linked to nucleotides. NtdA, a 3-oxo-glucose-6-phosphate aminotransferase^45^, and KdnA, a 8-amino-3,8-dideoxy-alpha-D-*manno*-octulosonate aminotransferase^46^ are examples of non-nucleotide sugar aminotransferases. These enzymes harbour variations in active site that accommodate nucleotide group. For example, the supposed nucleotide binding pocket of NtdA is occupied by side chains of Leu70 and Tyr246^45^. In RbmB, a 2-deoxy-*scyllo*-inosose aminotransferase^47^, a longer loop protrudes Phe336 into the nucleotide binding pocket, which along with Gln190 obstructs the binding of nucleotide group. Collectively, these observations suggest the presence of non-generic family specific features which cannot be deduced using sequence-based approaches.

The fold space of polypeptides is finite^48–52^. Consequently, the same fold can accommodate multiple functions using diverse mechanisms such as point mutations, indels, recruitment of additional domains, oligomerization, etc.^53^. Several factors such as gene loss, enzyme promiscuity, biological networks, etc., are proposed to drive neo- and subfunctionalization^54–56^. Aminotransferases and dehydratases are examples of functional families wherein a few mutations lead to alterations in reaction specificity^19,20^. In the absence of adequate amount of experimental data that capture sequence changes to functional changes at the level of substrate and reaction specificity, automated function prediction algorithms that provide functional annotation using sequence similarity are likely to lead to incomplete or erroneous annotations at the level of substrate and reaction specificity. Such limitations are reflected in the inability of C3_NSAT_HMM_ and C4_NSAT_HMM_ to identify C3 and C4 aminotransferases from hits of NSAT_HMM_. Limitations of profile HMMs are overcome by combining family specific motif search approach. Thus, in an attempt to provide clues to reaction specificity among aminotransferases, we have underscored the necessity of manual curation in protein function prediction and suggested approaches to overcome limitations of automated annotation algorithms.

PLP-dependent enzymes host a wide range of catalytic reactions and are highly relevant biocatalysts^57^. Minor alterations in the active site leading to functional diversity are being routinely engineered for biotechnological applications^58,59^. With experimental validation, features highlighted in this study that contribute to reaction specificities in aminotransferases and its homologs may be useful in engineering efficient enzymes for the biosynthesis of natural products.

## Abbreviations

Aminotransferase: Nucleotide Sugar Aminotransferase
Dehydratase: Nucleotide Sugar Dehydratase
HMM: Hidden Markov Model
PWM: Position Weight Matrix
MSA: Multiple Sequence Alignment
SDR: Short-chain Dehydrogenase Reductase
ROC: Receiver Operator Characteristic

## Data availability

PDB codes used in this study can be accessed from RCSB Protein Data Bank.

## Conflicts of Interests

None declared.

## Author Contributions

PVB conceptualized this study; JS performed analysis; JS and PVB co-wrote the paper.

## Acknowledgements

The authors are thankful to Professor Prasenjit Bhaumik for providing inputs on structural analysis. Jaya Srivastava is thankful to the Council of Scientific and Industrial Research, Government of India, for a research fellowship [number 09/087/(0877)/2017-EMR-I].

## Footnotes

a we used hits of profile HMMs instead of only the experimentally characterized sequences for this purpose with the expectation that a larger number of input sequences may lead to higher sensitivity of motif search

## Notes

### Competing Interest Statement

The authors have declared no competing interest.

### Summary of Updates

The usage of nucleotide sugar aminotransferases and dehydratases as NSATs and NSDs has been replaced with aminotransferases and dehydratases, respectively. Text concerning residue conservation analysis in dehydratases has been moved to discussion as it does not denote key results of this manuscripts. Some supplementary figures and tables have been removed. Prediction pipeline developed in this manuscript is validated with aminotransferases of established pathways whose activity has not been experimentally characterized.

