## Supplementary file for "Clues to reaction specificity in PLP-dependent fold type I aminotransferases of monosaccharide biosynthesis"

**Contents**


| | |
|---|---|
| Table S1 | Experimentally characterized C3 aminotransferases and their substrates and products |
| Table S2 | Experimentally characterized C4 aminotransferases and their substrates and products |
| Table S3 | Analysis of the extent of conservation of residues implicated in distinguishing transamination from dehydration |
| Figure S1 | A composite schematic of the pathways for the biosynthesis of monosaccharides |
| Figure S2 | All-against-all pairwise sequence identity matrix |
| Figure S3 | ROC curve of $NSAT_{HMM}$ and $NSD_{HMM}$ |
| Figure S4 | Superimposed ligands of C3 and C4 aminotransferases |
| Figure S5 | Structure and density maps of ligands of 2FNU and 4ZAH |
| Figure S6 | Phylogenetic subtree of PLP-dependent fold type I enzymes |
| References | References cited in supplementary data |

**Table S1**: Experimentally characterized C3 aminotransferases and their substrates and products¶

| Gene name | Substrate | Product |
|---|---|---|
| QdtB<br>WlarG<br>TylB | 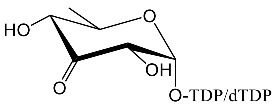 | 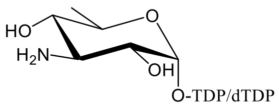 |
| FdtB<br>WlarG<br>RavAMT | 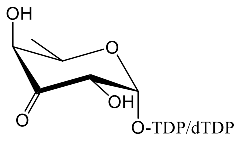 | 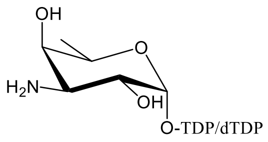 |
| DesV | 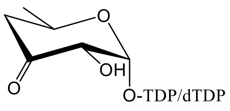 | 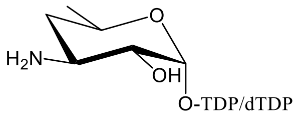 |
| EvaB<br>MegDII | 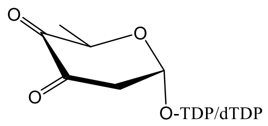 | 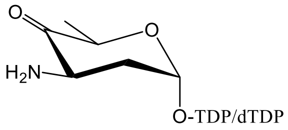 |
| WbpE | 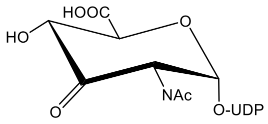 | 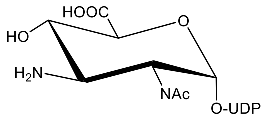 |

¶ Pyranose rings of substrates and products are shown in the $^4C_1$ conformation based on the configuration of the C5 atom (highest numbered chiral carbon in the pyranose ring). UniProt accession number, name and length of the protein, names of substrates and products, PDB ids of proteins for which 3D structure is known and PubMed IDs are given in the Supplementary data.xslx, worksheet: Exp and reviewed sequences.

**Table S2**: Experimentally characterized C4 aminotransferases and their substrates and products¶

| Gene name | Substrate | Product |
|---|---|---|
| DesI<br>VioA | 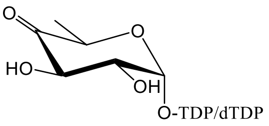 | 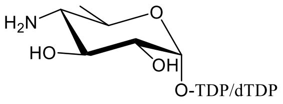 |
| WecE | 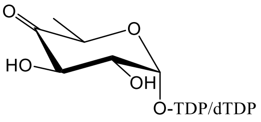 | 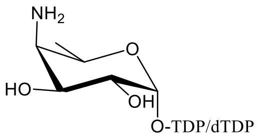 |
| LegC | 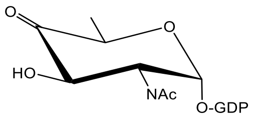 | 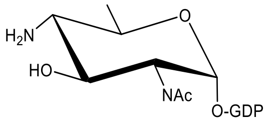 |
| WeeJ<br>PglC<br>PglE<br>EpsN | 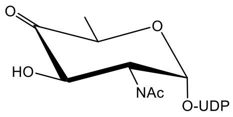 | 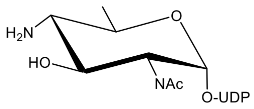 |
| Pat | 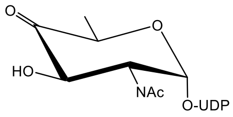 | 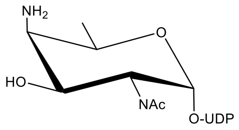 |
| PseC<br>Pam | 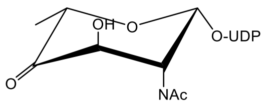 | 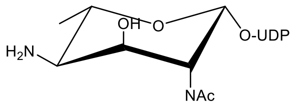 |
| Per<br>PerA<br>RfbE | 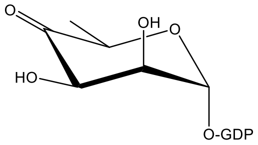 | 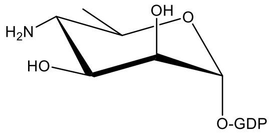 |
| ArnB | 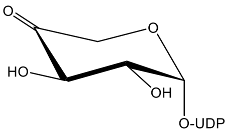 | 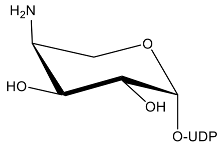 |
| SpnR | 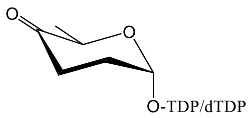 | 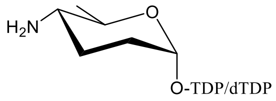 |

¶ Pyranose rings of substrates and products are shown in the ${}^4C_1$ or ${}^1C_4$ conformation based on the configuration of the C5 atom (highest numbered chiral carbon in the pyranose ring). UniProt accession number, name and length of the protein, names of substrates and products, PDB ids of proteins for which 3D structure is known and PubMed IDs are given in the Supplementary data.xslx, worksheet: Exp and reviewed sequences.

**Table S3:** Analysis of the extent of conservation of residues implicated in distinguishing transamination from dehydration

| Site | Site 1[¶] | | Site 2[¶] | | Site 3[‡] | | Site 4[¶‡@] | | Site 5[¶] | |
|---|---|---|---|---|---|---|---|---|---|---|
| Enzyme family (AT: aminotransferase; NSD: dehydratase) | AT | NSD | AT | NSD | AT | NSD | AT | NSD | AT | NSD |
| Residue | His | Asp[§] | His[£] | Tyr | Asn[†] | Ala | Lys | His | His[£] | Phe |
| Position[$] | 163 | 194 | 185 | 217 | 185 | 219 | 188 | 220 | 297 | 345 |
| C3_hits * (12811 sequences) | Gln: 12712<br>His: 41<br>Asp: 0 | | Tyr: 7703<br>Phe: 4954<br>His: 80 | | Ser: 4529<br>Gly: 4112<br>Thr: 3397<br>Ala: 603<br>Asn: 4 | | Lys: 12799<br>His: 0 | | His: 9605<br>Ala: 1389<br>Asn: 1108<br>Phe: 0 | |
| C4_hits * (12101 sequences) | His: 6868<br>Glu: 3602<br>Gln: 1482<br>Asp: 0 | | His: 6005<br>Asn: 2614<br>Tyr: 2462 | | Asn: 4126<br>Ile: 2363<br>Val: 2782<br>Thr: 2049<br>Ser: 71<br>Ala: 23 | | Lys: 12100<br>His: 0 | | His: 7469<br>Trp: 3414<br>Phe: 219 | |
| NSD_hits * (4018 sequences) | Asp: 3173<br>Glu: 844<br>His: 0 | | Tyr: 3249<br>Phe: 764<br>His: 5 | | Ala: 2303<br>Ser: 856<br>Pro: 822<br>Asn: 0 | | His: 4018<br>Lys: 0 | | Phe: 3783<br>His: 27 | |

[$] Positions in the primary structure to which the six sites correspond to and the residue at these positions correspond to *Salmonella Typhimurium* ArnB (*St*ArnB) for aminotransferase and *Yersinia pseudotuberculosis* E1 dehydratase for dehydratase.

* The number of sequences in which residue conservation analysis is done is given in parenthesis. For each site and enzyme family, the number of sequences in which the most frequently occurring residues / the residue in the reference proteins (mentioned in the column heading) occur are shown.

[¶] Data for Sites 1, 2, 4 and 5 are for *Yersinia pseudotuberculosis* E1 dehydratase. Wild type residues were replaced by residues at the homologous position in *St*ArnB, a C4 aminotransferase. The four site (D194H-Y217H-H220K-F345H) mutant showed aminotransferase activity [19].

[‡] Data for Sites 3 and 4 are for *Escherichia coli* GDP-4-keto 6-deoxy mannose 3-dehydratase (ColD). Wild type residues were replaced by residues at the homologous position in GDP-perosamine synthase (PerA), a C4 aminotransferase. The two site (S187A-H188K) mutant showed aminotransferase activity [20]. The residue at position 187 is Ala in *Yersinia pseudotuberculosis* E1 dehydratase.

[@] Lysine in this site forms Schiff's base with PLP. This lysine is replaced by histidine in dehydratases.

[§] Asp194 has been proposed to stabilize the ketimine intermediate through a salt bridge with 3-oxygen of the cofactor [65]. This residue is replaced by histidine in *St*ArnB.

[£] aminotransferases have histidine at Sites 2 and 5; these are not conserved in dehydratases. The role of these two histidines in transamination is not known.

[†] Asn185 in PerA H-bonds with the pyranose ring oxygen [63] and this H-bond is absent in ColD.

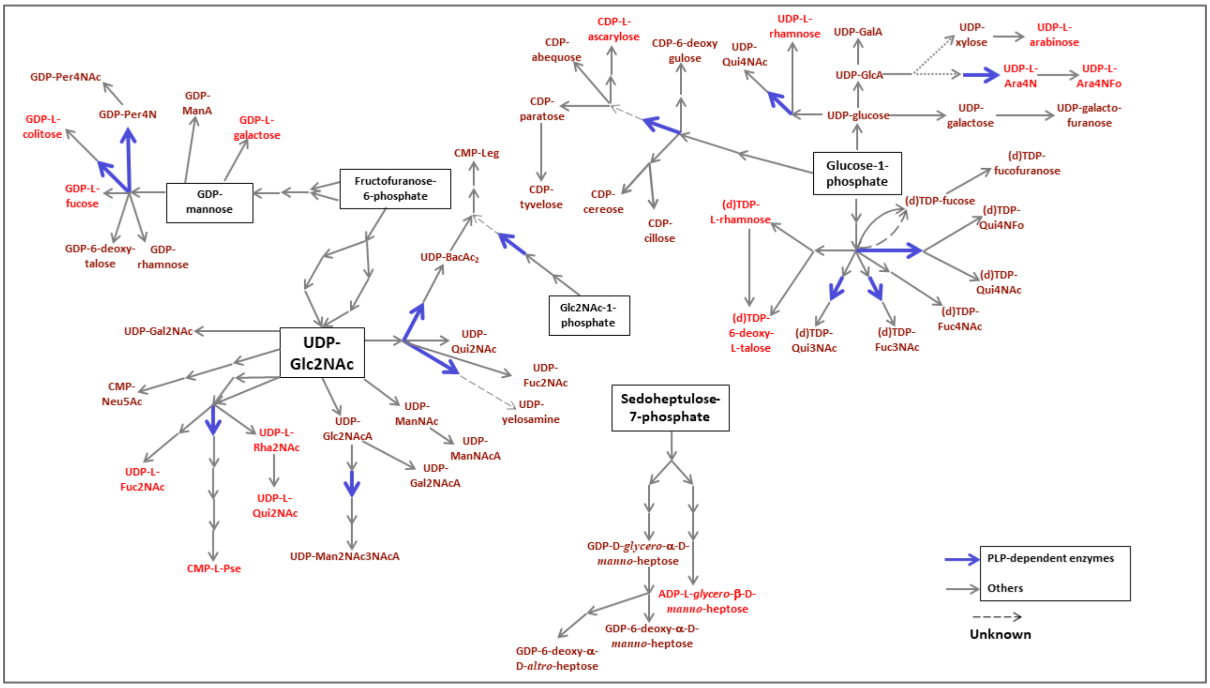


**Figure S1**: A composite schematic of the pathways for the biosynthesis of monosaccharides. Only the precursors (enclosed in a box) and end products of the pathways are mentioned by name. Products represented in maroon are D-sugars while those in red are L-sugars. Each arrow represents a step of a multistep pathway. Reactions catalyzed by nucleotide sugar aminotransferases are highlighted by blue arrows. Dashed arrows are used when the functional family to which the enzyme catalyzing the reaction is not clear. (d)TDP-linked sugars can be either dTDP- or TDP-linked; for the sake of simplicity, such sugars are mentioned as TDP-linked in this manuscript. This composite schematic is also provided in [1] but as a collage with names of enzyme families.

| | Q9F8 | Q9HZ | O528 | Q541 | D1H0 | Q9ZG | Q6TF | Q6T1 | Q9AL | P278 | Q814 | Q6U1 | Q9XC | Q9AL | Q9ZG | D6HA | B0V6 | Q795 | Q0P9 | Q0P8 | G8UH | Q8ZN | P776 | A0A1 | Q3ES | O251 | Q0P8 | Q9A9 | O683 | Q7DB | Q069 | B1B4 | Q9F1 | G4WJ | Q9AL | P263 | Q573 |
|---|---|---|---|---|---|---|---|---|---|---|---|---|---|---|---|---|---|---|---|---|---|---|---|---|---|---|---|---|---|---|---|---|---|---|---|---|---|
| Q9F837 | 100 | 36 | 71 | 43 | 45 | 46 | 39 | 40 | 36 | 29 | 24 | 24 | 22 | 30 | 31 | 29 | 29 | 28 | 24 | 22 | 24 | 31 | 30 | 28 | 29 | 22 | 22 | 35 | 29 | 29 | 28 | 23 | 23 | 22 | 27 | 28 | 28 |
| Q9HZ76 | 36 | 100 | 34 | 38 | 41 | 41 | 38 | 42 | 37 | 29 | 29 | 30 | 28 | 30 | 30 | 29 | 29 | 29 | 24 | 27 | 25 | 34 | 33 | 27 | 32 | 24 | 27 | 39 | 31 | 27 | 26 | 24 | 24 | 25 | 22 | 25 | 26 |
| O52807 | 71 | 34 | 100 | 41 | 43 | 45 | 37 | 39 | 34 | 29 | 27 | 23 | 25 | 32 | 31 | 30 | 30 | 27 | 24 | 22 | 20 | 30 | 29 | 29 | 31 | 24 | 23 | 34 | 30 | 28 | 27 | 23 | 23 | 22 | 24 | 26 | 26 |
| Q54142 | 43 | 38 | 41 | 100 | 58 | 58 | 44 | 48 | 38 | 29 | 26 | 24 | 25 | 31 | 33 | 30 | 30 | 27 | 23 | 22 | 23 | 35 | 35 | 29 | 29 | 24 | 25 | 37 | 31 | 31 | 27 | 21 | 21 | 24 | 28 | 23 | 23 |
| D1H0I5 | 45 | 41 | 43 | 58 | 100 | 59 | 44 | 49 | 39 | 29 | 25 | 24 | 24 | 32 | 35 | 29 | 29 | 26 | 22 | 23 | 24 | 35 | 34 | 32 | 31 | 23 | 25 | 35 | 33 | 29 | 30 | 24 | 24 | 24 | 26 | 25 | 26 |
| Q9ZGH4 | 46 | 41 | 45 | 58 | 59 | 100 | 45 | 49 | 42 | 29 | 26 | 25 | 26 | 33 | 33 | 33 | 30 | 26 | 24 | 22 | 22 | 37 | 37 | 29 | 31 | 25 | 24 | 38 | 37 | 32 | 30 | 21 | 21 | 24 | 28 | 24 | 23 |
| Q6TFC4 | 39 | 38 | 37 | 44 | 44 | 45 | 100 | 54 | 55 | 30 | 27 | 32 | 28 | 29 | 27 | 32 | 31 | 29 | 29 | 27 | 27 | 31 | 33 | 30 | 36 | 26 | 30 | 38 | 32 | 37 | 30 | 25 | 25 | 28 | 28 | 30 | 29 |
| Q6T1W6 | 40 | 42 | 39 | 48 | 49 | 49 | 54 | 100 | 64 | 29 | 25 | 28 | 26 | 29 | 28 | 31 | 31 | 26 | 24 | 25 | 22 | 34 | 36 | 32 | 36 | 26 | 25 | 37 | 33 | 37 | 30 | 25 | 25 | 26 | 26 | 28 | 28 |
| Q9ALS9 | 36 | 37 | 34 | 38 | 39 | 42 | 55 | 64 | 100 | 25 | 22 | 28 | 26 | 27 | 26 | 27 | 28 | 25 | 24 | 24 | 21 | 31 | 31 | 30 | 35 | 24 | 28 | 35 | 28 | 33 | 28 | 23 | 23 | 24 | 21 | 26 | 27 |
| P27833 | 29 | 29 | 29 | 29 | 29 | 29 | 30 | 29 | 25 | 100 | 28 | 28 | 25 | 28 | 25 | 25 | 25 | 29 | 25 | 26 | 26 | 32 | 32 | 27 | 28 | 23 | 25 | 32 | 28 | 30 | 27 | 19 | 19 | 20 | 28 | 30 | 30 |
| Q814Z4 | 24 | 29 | 27 | 26 | 25 | 26 | 27 | 25 | 22 | 28 | 100 | 28 | 27 | 27 | 23 | 27 | 26 | 27 | 26 | 23 | 22 | 32 | 33 | 31 | 32 | 28 | 33 | 31 | 28 | 31 | 27 | 23 | 23 | 25 | 20 | 21 | 21 |
| Q6U1I3 | 24 | 30 | 23 | 24 | 24 | 25 | 32 | 28 | 28 | 28 | 28 | 100 | 53 | 27 | 30 | 26 | 26 | 24 | 23 | 26 | 22 | 28 | 27 | 28 | 30 | 30 | 30 | 30 | 26 | 32 | 28 | 22 | 22 | 23 | 27 | 29 | 28 |
| Q9XCW4 | 22 | 28 | 25 | 25 | 24 | 26 | 28 | 26 | 26 | 25 | 27 | 53 | 100 | 27 | 34 | 22 | 23 | 25 | 23 | 24 | 23 | 29 | 29 | 27 | 31 | 32 | 28 | 29 | 27 | 32 | 30 | 20 | 20 | 21 | 24 | 26 | 25 |
| Q9ALN9 | 30 | 30 | 32 | 31 | 32 | 33 | 29 | 29 | 27 | 28 | 27 | 27 | 27 | 100 | 30 | 28 | 27 | 30 | 28 | 26 | 26 | 31 | 33 | 26 | 27 | 28 | 23 | 32 | 29 | 31 | 28 | 23 | 23 | 25 | 28 | 24 | 24 |
| Q9ZGH0 | 31 | 30 | 31 | 33 | 35 | 33 | 27 | 28 | 26 | 25 | 23 | 30 | 34 | 30 | 100 | 28 | 25 | 25 | 23 | 23 | 26 | 33 | 32 | 27 | 26 | 25 | 25 | 35 | 30 | 28 | 30 | 23 | 23 | 21 | 26 | 22 | 23 |
| D6HAT9 | 29 | 29 | 30 | 30 | 29 | 33 | 32 | 31 | 27 | 25 | 27 | 26 | 22 | 28 | 28 | 100 | 68 | 26 | 25 | 24 | 26 | 29 | 27 | 29 | 30 | 24 | 25 | 35 | 31 | 32 | 29 | 24 | 24 | 23 | 25 | 23 | 24 |
| B0V6J6 | 29 | 29 | 30 | 30 | 29 | 30 | 31 | 31 | 28 | 25 | 26 | 26 | 23 | 27 | 25 | 68 | 100 | 26 | 24 | 24 | 26 | 30 | 28 | 27 | 28 | 24 | 24 | 34 | 29 | 34 | 30 | 23 | 23 | 22 | 22 | 22 | 24 |
| Q795J3 | 28 | 29 | 27 | 27 | 26 | 26 | 29 | 26 | 25 | 29 | 27 | 24 | 25 | 30 | 25 | 26 | 26 | 100 | 48 | 38 | 36 | 31 | 32 | 27 | 30 | 26 | 30 | 36 | 34 | 36 | 35 | 24 | 24 | 25 | 24 | 26 | 27 |
| Q0P9D3 | 24 | 24 | 24 | 23 | 22 | 24 | 29 | 24 | 24 | 25 | 26 | 23 | 23 | 28 | 23 | 25 | 24 | 48 | 100 | 39 | 39 | 26 | 27 | 24 | 29 | 24 | 29 | 32 | 32 | 33 | 33 | 24 | 24 | 24 | 23 | 25 | 25 |
| Q0P8T7 | 22 | 27 | 22 | 22 | 23 | 22 | 27 | 25 | 24 | 26 | 23 | 26 | 24 | 26 | 23 | 24 | 24 | 38 | 39 | 100 | 48 | 26 | 29 | 22 | 27 | 27 | 28 | 32 | 33 | 35 | 34 | 23 | 23 | 22 | 21 | 24 | 25 |
| G8UHQ6 | 24 | 25 | 20 | 23 | 24 | 22 | 27 | 22 | 21 | 26 | 22 | 22 | 23 | 26 | 26 | 26 | 26 | 36 | 39 | 48 | 100 | 27 | 28 | 25 | 26 | 23 | 26 | 35 | 37 | 37 | 35 | 22 | 22 | 22 | 24 | 24 | 25 |
| Q8ZNF3 | 31 | 34 | 30 | 35 | 35 | 37 | 31 | 34 | 31 | 32 | 32 | 28 | 29 | 31 | 33 | 29 | 30 | 31 | 26 | 26 | 27 | 100 | 82 | 32 | 36 | 26 | 29 | 35 | 34 | 35 | 31 | 24 | 24 | 25 | 27 | 27 | 27 |
| P77690 | 30 | 33 | 29 | 35 | 34 | 37 | 33 | 36 | 31 | 32 | 33 | 27 | 29 | 33 | 32 | 27 | 28 | 32 | 27 | 29 | 28 | 82 | 100 | 31 | 34 | 28 | 28 | 35 | 34 | 34 | 29 | 24 | 24 | 25 | 28 | 29 | 28 |
| A0A1D3U | 28 | 27 | 29 | 29 | 32 | 29 | 30 | 32 | 30 | 27 | 31 | 28 | 27 | 26 | 27 | 29 | 27 | 27 | 24 | 22 | 25 | 32 | 31 | 100 | 47 | 33 | 36 | 32 | 30 | 34 | 31 | 24 | 24 | 25 | 23 | 26 | 25 |
| Q3ESA3 | 29 | 32 | 31 | 29 | 31 | 31 | 36 | 36 | 35 | 28 | 32 | 30 | 31 | 27 | 26 | 30 | 28 | 30 | 29 | 27 | 26 | 36 | 34 | 47 | 100 | 35 | 40 | 37 | 32 | 37 | 31 | 27 | 27 | 27 | 28 | 28 | 28 |
| O25130 | 22 | 24 | 24 | 24 | 23 | 25 | 26 | 26 | 24 | 23 | 28 | 30 | 32 | 28 | 25 | 24 | 24 | 26 | 24 | 27 | 23 | 26 | 28 | 33 | 35 | 100 | 45 | 28 | 24 | 27 | 25 | 24 | 24 | 23 | 21 | 22 | 21 |
| Q0P8W3 | 22 | 27 | 23 | 25 | 25 | 24 | 30 | 25 | 28 | 25 | 33 | 30 | 28 | 23 | 25 | 25 | 24 | 30 | 29 | 28 | 26 | 29 | 28 | 36 | 40 | 45 | 100 | 28 | 25 | 26 | 25 | 24 | 24 | 25 | 22 | 26 | 25 |
| Q9A9H3 | 35 | 39 | 34 | 37 | 35 | 38 | 38 | 37 | 35 | 32 | 31 | 30 | 29 | 32 | 35 | 35 | 34 | 36 | 32 | 32 | 35 | 35 | 35 | 32 | 37 | 28 | 28 | 100 | 46 | 46 | 41 | 26 | 26 | 27 | 29 | 29 | 29 |
| O68392 | 29 | 31 | 30 | 31 | 33 | 37 | 32 | 33 | 28 | 28 | 28 | 26 | 27 | 29 | 30 | 31 | 29 | 34 | 32 | 33 | 37 | 34 | 34 | 30 | 32 | 24 | 25 | 46 | 100 | 52 | 52 | 26 | 26 | 25 | 30 | 28 | 27 |
| Q7DBF3 | 29 | 27 | 28 | 31 | 29 | 32 | 37 | 37 | 33 | 30 | 31 | 32 | 32 | 31 | 28 | 32 | 34 | 36 | 33 | 35 | 37 | 35 | 34 | 34 | 37 | 27 | 26 | 46 | 52 | 100 | 54 | 27 | 27 | 28 | 28 | 30 | 30 |
| Q06953 | 28 | 26 | 27 | 27 | 30 | 30 | 30 | 30 | 28 | 27 | 27 | 28 | 30 | 28 | 30 | 29 | 30 | 35 | 33 | 34 | 35 | 31 | 29 | 31 | 31 | 25 | 25 | 41 | 52 | 54 | 100 | 26 | 26 | 26 | 28 | 27 | 27 |
| B1B4V9 | 23 | 24 | 23 | 21 | 24 | 21 | 25 | 25 | 23 | 19 | 23 | 22 | 20 | 23 | 23 | 24 | 23 | 24 | 24 | 23 | 22 | 24 | 24 | 24 | 27 | 24 | 24 | 26 | 26 | 27 | 26 | 100 | 100 | 74 | 31 | 29 | 29 |
| Q9F118 | 23 | 24 | 23 | 21 | 24 | 21 | 25 | 25 | 23 | 19 | 23 | 22 | 20 | 23 | 23 | 24 | 23 | 24 | 24 | 23 | 22 | 24 | 24 | 24 | 27 | 24 | 24 | 26 | 26 | 27 | 26 | 100 | 100 | 74 | 31 | 29 | 29 |
| G4WJD4 | 22 | 25 | 22 | 24 | 24 | 24 | 28 | 26 | 24 | 20 | 25 | 23 | 21 | 25 | 21 | 23 | 22 | 25 | 24 | 22 | 22 | 25 | 25 | 25 | 27 | 23 | 25 | 27 | 25 | 28 | 26 | 74 | 74 | 100 | 30 | 30 | 30 |
| Q9ALN8 | 27 | 22 | 24 | 28 | 26 | 28 | 28 | 26 | 21 | 28 | 20 | 27 | 24 | 28 | 26 | 25 | 22 | 24 | 23 | 21 | 24 | 27 | 28 | 23 | 28 | 21 | 22 | 29 | 30 | 28 | 28 | 31 | 31 | 30 | 100 | 52 | 51 |
| P26398 | 28 | 25 | 26 | 23 | 25 | 24 | 30 | 28 | 26 | 30 | 21 | 29 | 26 | 24 | 22 | 23 | 22 | 26 | 25 | 24 | 24 | 27 | 29 | 26 | 28 | 22 | 26 | 29 | 28 | 30 | 27 | 29 | 29 | 30 | 52 | 100 | 87 |
| Q57323 | 28 | 26 | 26 | 23 | 26 | 23 | 29 | 28 | 27 | 30 | 21 | 28 | 25 | 24 | 23 | 24 | 24 | 27 | 25 | 25 | 25 | 27 | 28 | 25 | 28 | 21 | 25 | 29 | 27 | 30 | 27 | 29 | 29 | 30 | 51 | 87 | 100 |

C3-NSAT Axial ($^4C_1$)
C3-NSAT Equatorial ($^4C_1$)
C4-NSAT Axial ($^4C_1$)
C4-NSAT Equatorial ($^4C_1$)
NSDH

**Figure S2:** All-against-all pairwise sequence identity matrix. Proteins are identified by their UniProt accession number and are arranged in the same order in both rows and columns. Cells of the matrix are coloured using MS-Excel's conditional formatting option to visually highlight sequence pairs that share "higher" similarity with each other.

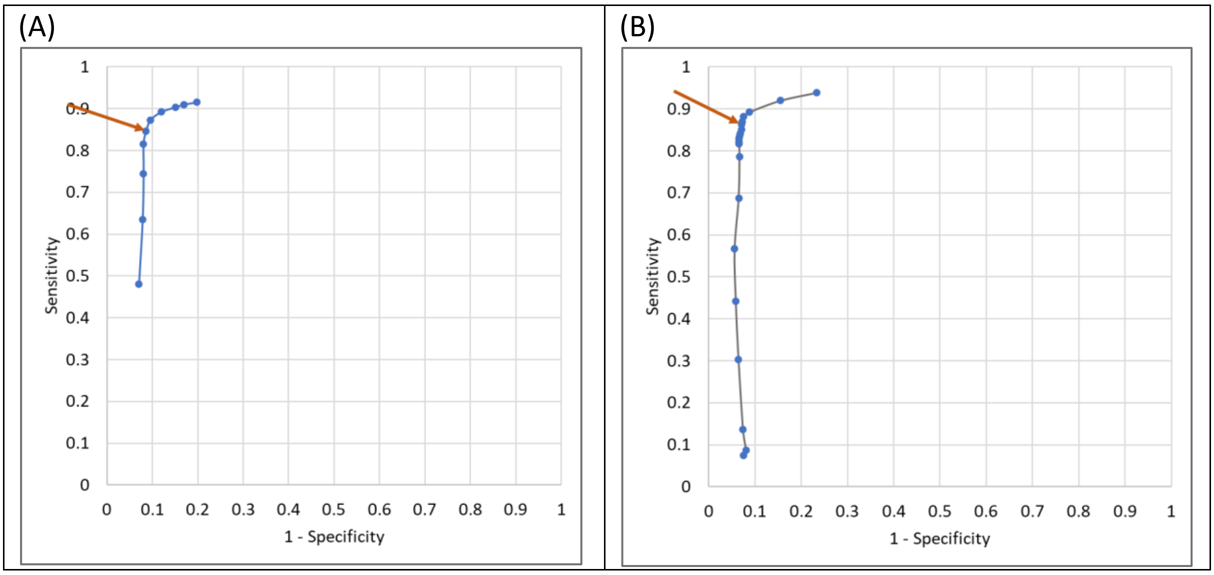


**Figure S3:** ROC curve generated from hits of (A) $NSAT_{HMM}$ and (B) $NSD_{HMM}$. Bit score threshold was chosen corresponding to the data point indicated by arrow.

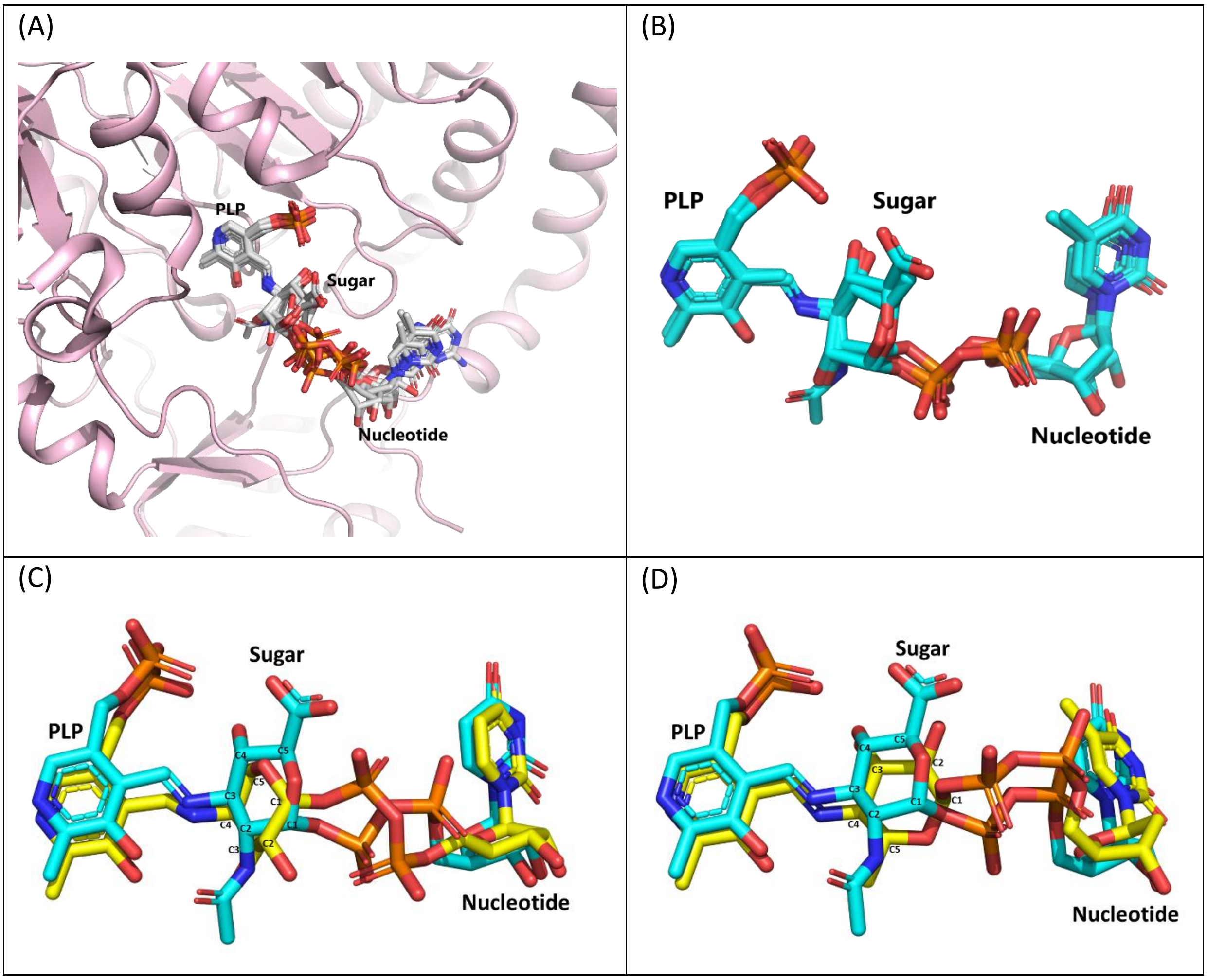


**Figure S4:** (A) Superposition ligands observed by structural superposition Cα backbone of all structures given in Table S1. (B) Comparison of substrate conformation of C3 aminotransferases WbpE (3NYT), WlarG (5U21) and QdtB (3FRK). (C) Comparison of substrate conformation in WbpE (blue) and ArnB (4OCA, yellow) depicting a shift in pyranose ring. (D) Comparison of substrate conformation in WbpE (yellow) and DesI (2PO3, yellow) depicting a flip in pyranose ring.

(A)

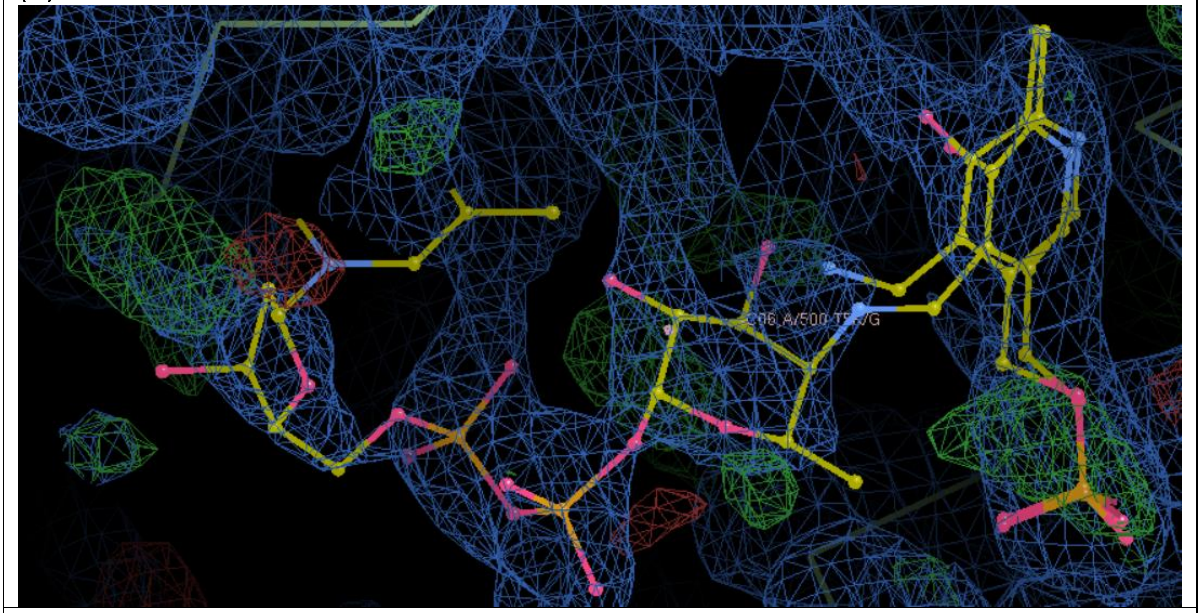

(B)

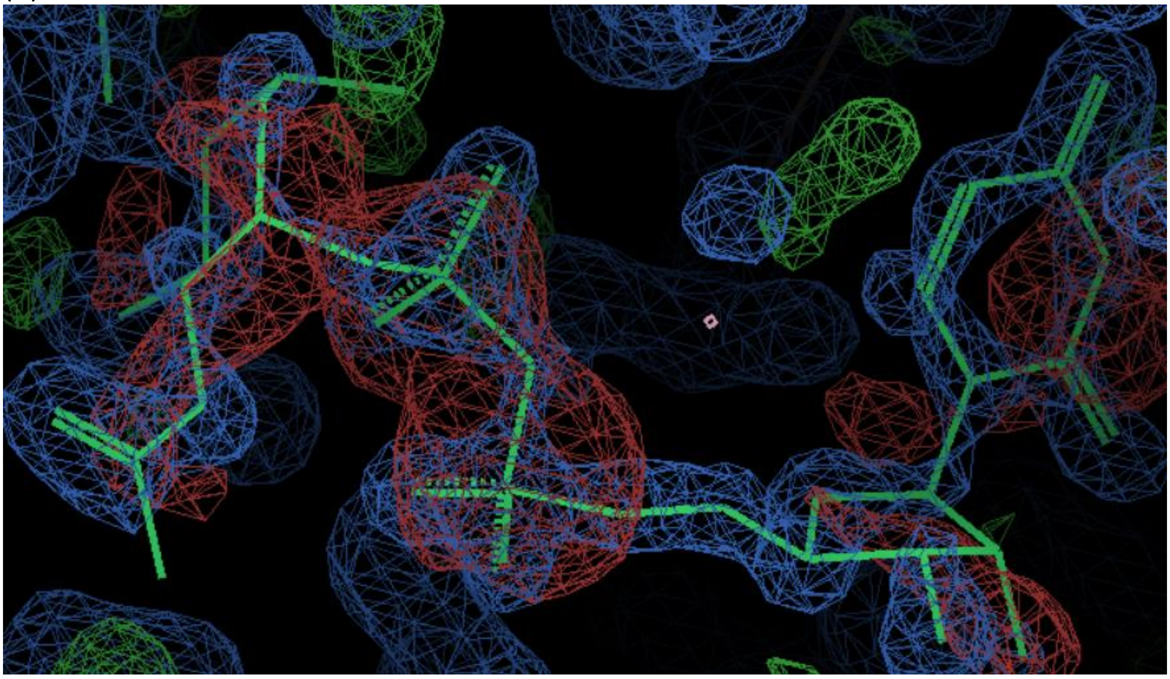

**Figure S5**: Screenshots of ligand structures fit in their density maps for A) 4ZAH and B)2FNU

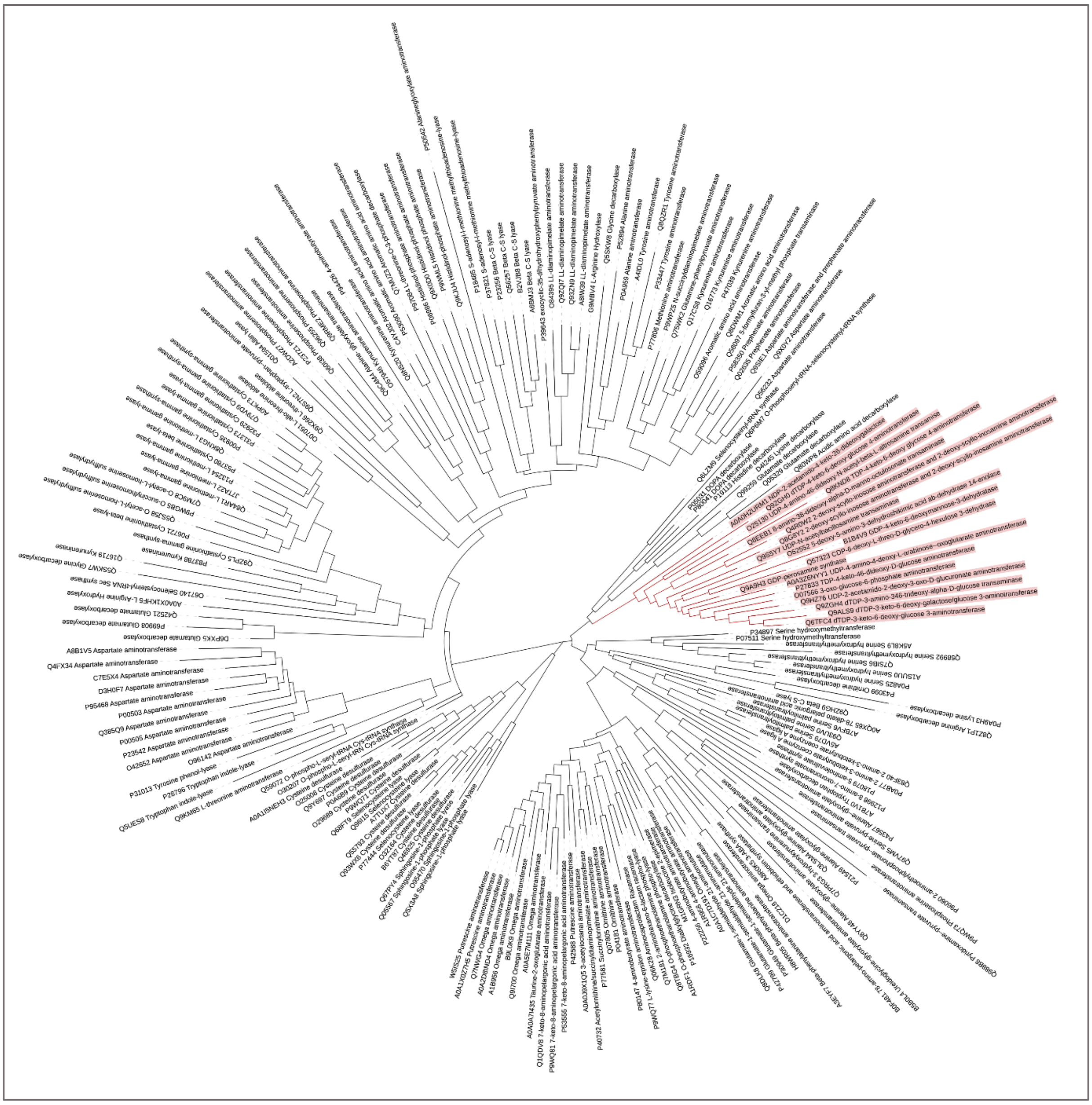

**Figure S6:** Phylogenetic tree of fold type I PLP-dependent enzymes. The clade containing aminotransferases and its homologs are highlighted in red.
